# A simple and reliable method for claustrum localization across age in mice

**DOI:** 10.1101/2023.10.31.564789

**Authors:** Tarek Shaker, Gwyneth J. Dagpa, Vanessa Cattaud, Brian A. Marriott, Mariam Sultan, Mohammed Almokdad, Jesse Jackson

## Abstract

The anatomical organization of the rodent claustrum remains obscure due to lack of clear borders that distinguish it from neighboring forebrain structures. Defining what constitutes the claustrum is imperative for elucidating its functions. Methods based on gene/protein expression or transgenic mice have been used to spatially outline the claustrum but often report incomplete labeling and/or lack of specificity during certain neurodevelopmental timepoints. To reliably identify claustrum cells in mice, we propose a simple immunolabelling method that juxtaposes the expression pattern of claustrum-enriched and cortical-enriched markers. We determined that claustrum cells immunoreactive for the claustrum-enriched markers Nurr1 and Nr2f2 are devoid of the cortical marker Tle4, which allowed us to differentiate the claustrum from adjoining cortical cells. Using retrograde tracing, we verified that nearly all claustrum projection neurons lack Tle4 but expressed Nurr1/Nr2f2 markers to different degrees. At neonatal stages between 7 and 21 days, claustrum projection neurons were identified by their Nurr1-postive/Tle4-negative expression profile, a time-period when techniques used to localize the claustrum in adult mice are ineffective. Finally, exposure to environmental novelty enhanced the expression of the neuronal activation marker cFos in the claustrum region. Notably, cFos labeling was mainly restricted to Nurr1-positive cells and nearly absent from Tle4-positive cells, thus corroborating previous work reporting novelty-induced claustrum activation. Taken together, this method will aid in studying the claustrum during postnatal development and may improve histological and functional studies where other approaches are not amenable.

## 2. INTRODUCTION

There is a longstanding concept in science that function can be informed by structure. The claustrum, which is a thin aggregate of neurons in the forebrain, forms extensive reciprocal connections with many cortical and subcortical regions (1–6). This broad networking feature of the claustrum is a main contributing factor as to why the claustrum is implicated in a wide range of functions that include consciousness processing, attention, and coordination of signal processing within the brain [reviewed in (6)]. However, up to this point, the exact role of the claustrum in the functions assigned to it remains poorly understood. One main challenge of elucidating the mechanisms underlying claustrum functional properties is that there is no consensus on what constitutes the claustrum (7–9). The definition of the claustrum is more debatable in rodents than in primates (10). This is mostly because the primate external capsule separates the claustrum from the medially located striatum and the extreme capsule separates the claustrum from the insula (10). However, because rodents lack the extreme capsule, there are no anatomical borders that distinguish the claustrum from the insula (10). Therefore, there is a pressing need for anatomical delineation of the claustrum, especially in rodents.

Due to a lack of definitive anatomical landmarks in rodents, the molecular profile of genes that preferentially label claustrum cells can be in theory used to locate the claustrum. Although many genes such as Nr4a2 (also known as Nurr1), Nr2f2, Ntng2, Gnb4 and Lxn, are densely expressed in claustrum neurons, these genes are also sparsely expressed in the spatially adjacent sensory cortex, insula, and dorsal endopiriform nucleus (3,11–13). Another common approach to locate the claustrum is to employ retrograde labelling of claustrum projection neurons. Injection of retrograde tracers into cortical regions that receive claustrum inputs labels distinct cell populations organized in modules across the dorsoventral axes of the claustrum (14–16). However, the labelling efficiency of retrograde tracing from a single cortical region is 50-60%, leading to incomplete labelling of the claustrocortical cells, and in some cases neighboring structures such as the insula and dorsal endopiriform are also labelled by retrograde tracing (15,17). Alternatively, some studies have taken advantage of the strong neuropil labelling exhibited by parvalbumin (PV) expressing interneurons in the claustrum core relative to the shell region in order to locate the core domain of the claustrum (3,15,18–20). However, labelling claustrum interneurons rather claustrum principal neurons, i.e. projection neurons, fails to define claustrum borders or to segregate claustrum cells from their surroundings. PV labelling is weak or absent at young ages which precludes the use of this method for developmental studies (20). Therefore, each of the aforementioned strategies used for defining the spatial location of the claustrum has strengths and tradeoffs depending on the experimental goal. Our goal here was to provide a complimentary method to establish the boundaries of the claustrum and discriminate it from nearby structures in mice.

We and others have previously described the expression profile of genes that are devoid from the claustrum and yet highly enriched in surrounding cortical structures, e.g. Tle4, Nnat and Ctgf (3,8,12,21–25). Thus, we set out to find candidate cortical-enriched markers that do not colocalize with claustrum-enriched markers, i.e. claustrum-devoid cortical markers. Our aim was to exploit the contrast between the labelling of claustrum-enriched and claustrum-devoid markers to highlight claustrum borders and to help identify claustrum projection neurons. In mice, we found that Tle4 is conspicuously absent from claustrum projection neurons as measured by retrograde labelling from the cortex. This pattern was consistent across different populations of claustrum projection neurons suggesting that Tle4 is indeed a claustrum-devoid marker. We also revealed a lack of colocalization between Tle4 and the claustrum-enriched markers Nurr1 and Nr2f2. Conversely, Tle4 was highly co-expressed with Nurr1 in cortical areas outside of the claustrum. Therefore, the combination of Tle4 and Nurr1 (or Nr2f2) labelling aids to better visualize the claustrum borders and to identify claustrum projection neurons. We also show that this approach demarcates the claustrum across early post-natal development where other methods are not effective. Finally, we show that environmental novelty induces cFos activation of Nurr1-enriched/Tle4-devoid cells within the claustrum, in line with previous data (26,27). Together, we demonstrate that the expression pattern of Tle4 combined with claustrum-enriched markers is a reliable method for anatomical demarcation of claustrum neurons across early postnatal and adult ages alike.

## 3. MATERIALS AND METHODS

### 3.1. Experimental Animals

Naive mice were maintained on a C57BL/6 background. For the majority of experiments, adult mice were used between 70 and 100 days old. For developmental experiments, neonatal mice between day 0 and 21 (±1 day) and young adult mice 49 ± 1 days old were used. Mice were group-housed under standard pathogen-free conditions, in a temperature-controlled environment and 12 h light/dark cycle, and with ad libitum access to water and food. Male and female mice were included in the study (see figure legends and **Supplementary Tables 1, 3, 4** for details on the sex of mice used for each experiment).

### 3.2 Stereotaxic injection of viral vectors

Adeno-associated viruses (AAVs) obtained from Addgene (MA, USA) were injected in different cortical regions. Mice at postnatal day (P) 0, P7, P14 and P49 were injected in the anterior cingulate cortex (ACC) with retrograde pAAV-CAG-tdTomato (59462-AAVrg), and mice >P70 were injected in the ACC, lateral entorhinal cortex (LEC), primary motor cortex (MOp) and retrosplenial cortex (RSC) with retrograde pAAV-CAG-GFP (37825-AAVrg). All injections were performed unilaterally in the left hemisphere. For injections in neonatal mice, the volume was optimized as previously described in (28) to ensure viral infusion selectivity to the target cortical region without spreading to adjacent regions. Injection volume for mice at P0 was 75 nl, at P7 was 100 nl and at P14 was 125 nl. For mice at P49 and >P70, the volume was 200 nl.

#### 3.2.1. Neonatal mice at P0

Newborn pups at P0 were collected from their home cage and prepared for surgery by cryoanesthesia. Following cessation of movement, the pup was positioned in a homemade head holder fitted to the stereotaxic apparatus. Injection site coordinates were determined relative to the intersection of the transverse sinus and the superior sagittal sinus. Stereotaxic coordinates were A/P: +2.35, M/L: +0.20, D/V: -0.20. A glass pipette (pulled at ∼10 µm in diameter) backfilled with mineral oil and loaded with tracers was slowly lowered to penetrate the skin until the pipette tip contacts the surface of the skull. D-V coordinates were set at zero. Then, the pipette was lowered to the target site, and AAVs were injected at a rate of 50 nl/min. Once 75 nl of AAVs was injected, the pipette was left in position for one minute before bringing it up slowly to half D-V depth. After one minute, the pipette was slowly withdrawn from the pup’s head. Thereafter, the pup was allowed to recover on a heating pad until it regained normal color and resumed movement before it was returned to the dam.

#### 3.2.1. Pre-weaned postnatal mice at P7 and P14

For pups between P7 and P14, carprofen (2 mg/kg) was administered subcutaneously 5-10 minutes before the surgery. Anesthesia was induced at 3.5-4.0% isoflurane and thereafter maintained at 1.5–2.0% during surgery, with O_2_ flow at ∼0.6L/min. The head was fixed in a homemade head holder customized for juvenile pups while the mouse was resting on a heating pad to maintain the body temperature at 37°C throughout the surgery. Mouse reflex was evaluated to assess the depth of anesthesia. Fur covering the scalp was removed with a fine trimmer. For P14 mice, eyes were covered with ointment to avoid their dryness. Under sterile conditions, the scalp was disinfected with 70% ethanol and betadine. Bupivacaine was applied under the scalp, and a small incision was made along the midline. The skull was exposed and leveled along A-P and M-L coordinates relative to the bregma. While the incision was open, the skull was kept moist with warm sterile 0.9% saline. A fine needle was used to poke a hole in the skull at injection site coordinates relative to bregma. Similar to above, a glass pipette backfilled with mineral oil and loaded with AAV was lowered to the target site. Stereotaxic coordinates for the ACC were the following: P7, A/P: +0.40, M/L: +0.25, D/V: -0.30; P14, A/P: +0.65, M/L: +0.30, D/V: -0.40. DV coordinates were measured from brain surface. Once the full amount of AAVs was injected, the pipette was left in position for ∼5 minutes before being slowly withdrawn from the pup’s head. The skin was resealed with sutures and Vetbond tissue adhesive (3M, MN, USA). Following mouse recovery on a heating pad, it was returned to its home cage.

#### 3.2.3. Adult mice (P49 and >P70)

For adult mice, stereotaxic injections were performed as described above for P7 and P14 mice, with the following exceptions: Carprofen was administered in water (5 mg/kg) ad libitum 24 hr prior to surgery, and for 72 hr after surgery. The mouse was placed in a stereotaxic frame, and an ointment was applied to its eyes to avoid their dryness. A dental drill was used to make small openings on top of the injection site. After the injection is complete, the pipette was left in position for 10-12 minutes before withdrawal. Stereotaxic coordinates were the following: ACC, A/P: +1.70, M/L: +0.50, D/V: -0.70; LEC, A/P: -3.00, M/L: +4.20, D/V: -2.35; MOp, A/P: +1.00, M/L: +1.70, D/V: -0.80; RSC, A/P: -2.00, M/L: +0.50, D/V: -0.50.

### 3.3. Open field behavioural test

Mice at 2-3 months were used for open field (OF) test. All behavioral experiments were conducted between 9 A.M. and 3 P.M. Mice were first habituated to the handler for two days before running OF. At the beginning of each session, mice were habituated to the room for 30 minutes before being handled by the experimenter. Mice were handled for two 5-minute sessions twice a day over a 2-day period under a reverse 12-hour light-dark cycle. Each session was spaced out by 3 hours. OF apparatus (25 cm x 25cm opaque walled box with open top) was sterilized using 50% ethanol and water prior to behavioral testing and in between mice. Mice that were randomly selected to be placed in OF were gently placed into the middle of the apparatus and allowed to explore for 10 minutes, before being returned to their home cage. Littermates that remained in the home cage throughout the OF experiment, i.e. naive mice, were deemed as controls. Afterwards, all mice were left undisturbed for 60-90 minutes before tissue collection. All experiments were recorded using overhead camera (20 frames/s) connected to FlyCap2 software (Teledyne FLIR, OR, USA).

### 3.4. Tissue collection and processing

Mice were deeply anesthetized and transcardially perfused with 4% paraformaldehyde (PFA) in 1x phosphate buffer solution (PBS), pH 7.4. Brains were extracted and fixed overnight in 4% PFA at 4℃, followed by washing three times with 1x PBS for 5 min, and then stored in 1x PBS. Depending on the experiment, mice were perfused either 7 or 14 days after stereotaxic injection of AAVs. For tissue sectioning, brains were submerged in pre-warmed 2% liquid agarose dissolved in 1x PBS, and brain-containing agarose blocks were mounted with superglue. Coronal brain slices were cut at 50 μm with a vibratome (5100mz; Campden Instruments, UK). Slices were collected starting at the level of the anterior insula, along the A-P axis, up until the level of the ventral hippocampus.

### 3.5. Immunohistochemistry

Slices were permeabilized by 10 min washing in 0.3% Triton X-100 in 1x PBS. Next, slices were incubated in a blocking solution, containing 3% bovine serum albumin and 0.3% Triton X-100 in 1x PBS, for 2 h at room temperature, and then incubated overnight at 4℃ in primary antibodies diluted in the same blocking solution. The following primary antibodies were used at the indicated dilutions: goat anti-Nurr1 (AF2156, R&D Systems; 1:250), rabbit anti-Nr2f2 (ab211776, Abcam; 1:250), mouse anti-Tle4 (sc365406, Santa Cruz Biotechnology; 1:250), goat anti-PV (PVG213, Swant; 1:2000), rat anti-SST (MAB354, Millipore; 1:250), rabbit anti-cFos (2250S, Cell Signaling; 1:1,000). The next day, slices were washed three times in 1x PBS for 10 min, then incubated with secondary antibodies diluted in blocking solution for 2 h at room temperature, before washing the slices again three times with 1x PBS for 10 min. The following secondary antibodies (ThermoFisher Scientific, MA, USA) were used at 1:500 dilution: Donkey anti-Goat IgG Alexa Fluor 488 (A-11055), Donkey anti-Goat IgG Alexa Fluor 555 (A-21432), Donkey anti-Rabbit IgG Alexa Fluor 555 (A-31572), Donkey anti-Mouse IgG Alexa Fluor 647 (A-31571), Chicken anti-Rat IgG Alexa Fluor 647 (A-21472). Finally, slices were mounted onto slides with Prolong Gold (P36930; ThermoFisher Scientific) and cover slipped.

### 3.6. Imaging

All images were taken on Leica TCS-SP5 and Leica SP8-STED confocal microscopes (Leica, Germany). Non-overlapping images from a single confocal plane were acquired with a 10x objective (0.3 NA for TSC-SP5 and 0.40 NA for SP8-STED), and 25x objective (0.5 NA for TSC-SP5 and 0.95 NA for SP8-STED). White light laser set at 488, 543, and 633 nm was used in a sequential order to eliminate any potential crosstalk of different channels. The acousto-optical emission filtering was set for Leica defaults of AF488, AF555 and AF647. Image acquisition settings were the following: pinhole 1 airy unit, scan speed 400 Hz unidirectional, format 1024 × 1024 pixels, z-step size 2 µm over 20 µm volume (Total of 11 images for each z-stack). Images were acquired using the same acquisition parameters for each fluorescent tracer/antibody labelling, with parameters adjusted to optimize brightness and contrast for each channel. Single-channel images were pseudocoloured to make data visualization more accessible to the readers. For each mouse, images were taken for 6 slices containing the claustrum, spanning the anterior, middle and posterior subdivisions, with 2 slices per subdivision separated by 100-200 µm. Only the hemisphere ipsilateral to injection site was imaged. All images were processed using FIJI (ImageJ, NIH, MD, USA) for analysis.

### 3.7. Analysis

Mice were included in this study only when the injection site was confirmed to be in the target cortical region. Quantification of confocal images was performed on maximum intensity projection of z-stacks. Using Matlab (Mathworks, MA, USA), the claustrum was demarcated based on cell labelling in the retrograde tracer channel, then cells were counted and cell location was registered in each imaging channel as previously described in (15). For spatial analysis in Figure 2, comparing spatial distribution between different markers for each image was achieved by manually plotting the perimeter of the zone enriched with labeled cells (>90%) in each imaging channel, resulting in a polygon that outlines the boundaries of the enriched zone. Single-channel images of the same slice that contain spatially registered polygons for two different markers were superimposed, and coordinates for the polygons were spatially registered simultaneously. Channel pairing combinations were GFP/Nr2f2 and GFP/Tle4. The centroids of the polygons were determined based on the coordinates of the vertices. Then, GFP/Nr2f2 or GFP/Tle4 polygon pairs from the same anteroposterior claustrum subdivision were overlaid to obtain the cumulative spatial distribution of that subdivision across all animals. This was performed while aligning the centroids of the polygons outlining the GFP zone, thus serving as a reference for polygon alignment. Statistics and bar graphs were produced using GraphPad Prism 8.0 (Dotmatics, MA, USA). Venn diagrams were generated by Matlab. Differences between three or more experimental groups were assessed with one-way ANOVA followed by Bonferroni post hoc comparison. Differences between two groups were assessed by unpaired t-test. Mean differences were considered to be significant at p < 0.05, with n being the number of mice analyzed. For spatial analysis in Figure 7, we generated spatial fluorescence intensity plots for retrograde tracing and cortical/claustrum markers by measuring the intensity in two axes: one orientated along the white matter parallel with the external capsule, and the other perpendicular to this axis. Spatial fluorescence profiles were calculated by taking the mean pixel intensity across a 0.1mm section in each axis. The intensity profiles were z-scored and realigned to be centered on the peak fluorescence of the retrograde labelled cells, thereby enabling averaging across brain sections and mice.

## 4. RESULTS

### 4.1. Coupling the expression profile of Tle4 with claustrum-enriched markers facilitates locating the claustrum

Currently available claustrum marker genes, like Nurr1 and Nr2f2 (**Supplementary Figure 1A**), are expressed, albeit sparsely, in the adjoining endopiriform nucleus, insula and gustatory-visceral cortex (3,11–13), resulting in lack of claustrum specificity for these markers. Thus, there is a need for marker genes that separate claustrum cells from adjacent cortical cells. We reasoned that anatomical delineation of the claustrum could be achieved by leveraging the combinatorial molecular profiles of claustrum-enriched and cortical-enriched genes.

In a recent transcriptomic study, it was reported that a number of genes distinguish projection neurons in the claustrum from nearby cortical neurons (25). We compared these findings with the Allen mouse brain in situ hybridization database (29) to identify candidate cortical-enriched genes that are devoid from the claustrum at early postnatal developmental stages and in adulthood (**Supplementary Figure 1B**). We noticed that the transcription factor Tle4 satisfied both conditions better than other candidates: 1) single-cell transcriptomic molecular analysis shows that Tle4 is absent in the claustrum (25), and 2) Spatial expression of Tle4 appears to be highly enriched in deep layers of the insula as well as in deep layers of the sensory cortex, while avoiding the claustrum, both at P4 and P56 (**Supplementary Figure 1B**). We identified an anti-Tle4 antibody that worked reliably for brain tissue derived from mice at different ages. Therefore, we set out to investigate whether Tle4 protein expression corroborates previous findings at the transcriptional level.

Topographical mapping of the claustrum has previously defined claustrum neurons projecting to the RSC as the center of the claustrum core, hereafter referred to as the central zone (15,30). Thus, for the first set of experiments, we used AAV retrograde fluorescent labelling of the RSC (Retro-RSC) as our reference to the claustrum. We performed unilateral injections of retro-AAV-CAG-GFP into the RSC of adult mice (>P70) and analyzed GFP expression in the claustrum 14 days post injection (**Figure 1A-C**). Several studies found Nurr1 to be a relatively selective marker for the claustrum at different ages and across different species, including rodents and primates (10,11,13,20). We found that Nurr1 immunolabelling showed a dense population of cells that appeared to be spatially aligned with the GFP-labeled retro-RSC central zone (**Figure 1C, D**). In contrast, Tle4 immunolabelling revealed that Tle4 was absent in the central zone, whereas outside of the central zone, Tle4 expression was noticeable in regions bordering the claustrum at all sides (**Figure 1E, F**). This suggests that while Nurr1 is a claustrum-enriched marker, Tle4 is a potential claustrum-devoid marker. Next, we quantified colocalization of RSC-projecting cells with Nurr1 and with Tle4 in the anterior, middle and posterior subdivisions of the claustrum. While a large proportion of retro-RSC-projecting cells co-expressed Nurr1 (>87%) (**Figure 1G, I, J, L, M, O**), very few cells co-expressed Tle4 (∼1%) (**Figure 1H, I, K, L, N, O**). Of note, these results were consistent in all claustrum subdivisions along the anteroposterior axis, and thus verify absence of Tle4 expression in projection neurons throughout the central zone of the claustrum.

**Figure 1:**
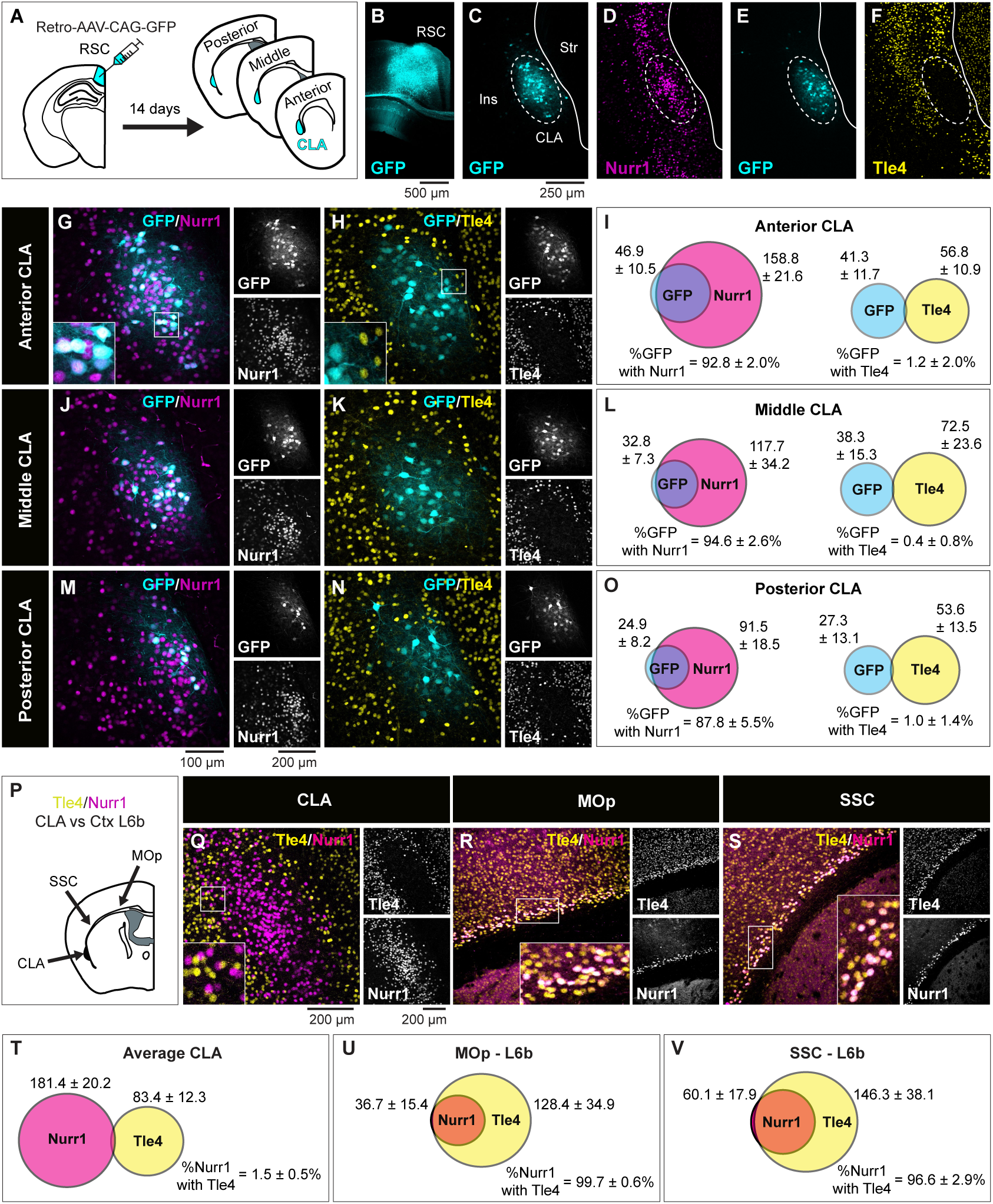
The expression pattern of Nurr1 and Tle4 in the claustrum region distinguishes claustrum projection neurons from their cortical counterparts in layer 5 and 6. **A:** Retrograde AAV-CAG-GFP was injected into the retrosplenial cortex (RSC), followed by tissue collection 14 days post injection. **B:** An example tracer injection site in the retrosplenial cortex**. C, E:** representative images of retrograde GFP labeling in the anterior claustrum **(D, F),** and representative images showing Nurr1 (**D**) and Tle4 (**F**) expression relative to GFP labeling. **C-D** are from the same slice, and **E-F** are from an adjacent slice of the same mouse. Dashed ellipses in **C-F** highlight GFP expression in the claustrum (CLA) without labeling the surrounding insula cortex (Ins) or striatum (str). **G-H:** Representative images showing the degree of colocalization with Nurr1 (**G**) or Tle4 labelled cells (**H**). Single channel images are shown on the right of each panel. **I**: Venn diagrams showing the mean cell counts for Nurr^+^/GFP^+^ and Tle4^+^/GFP^+^ cells in the anterior CLA. Values throughout indicate mean ± standard deviation (n = 6 mice, 3 male and 3 female). **J-L**: The same as **G-I** but for the middle CLA. **M-O**: The same as **G-I** for the posterior CLA. **P:** Nurr1 and Tle4 colocalization in the CLA region was compared to layer 6b (Ctx L6b) of the primary motor cortex (MOp) and the somatosensory cortex (SSC). **Q-S:** Representative images showing Nurr1 and Tle4 expression in the CLA region (**Q**), the MOp (**R**) and the SSC (**S**) (left, merged imaging channels; right, single channel images). **T-V:** Venn diagrams representing the mean number of cells expressing Nurr1 (magenta) and Tle4 (yellow), and colocalization between Nurr1 with Tle4 (n = 4 mice, 2 male and 2 female). The data in T are averaged across anterior, middle, and posterior CLA. Values in I, L, O, T, U, V represent mean ± standard deviation. Panels P-V: See **Supplementary Table 1** for details on mouse sex.

**Figure 2:**
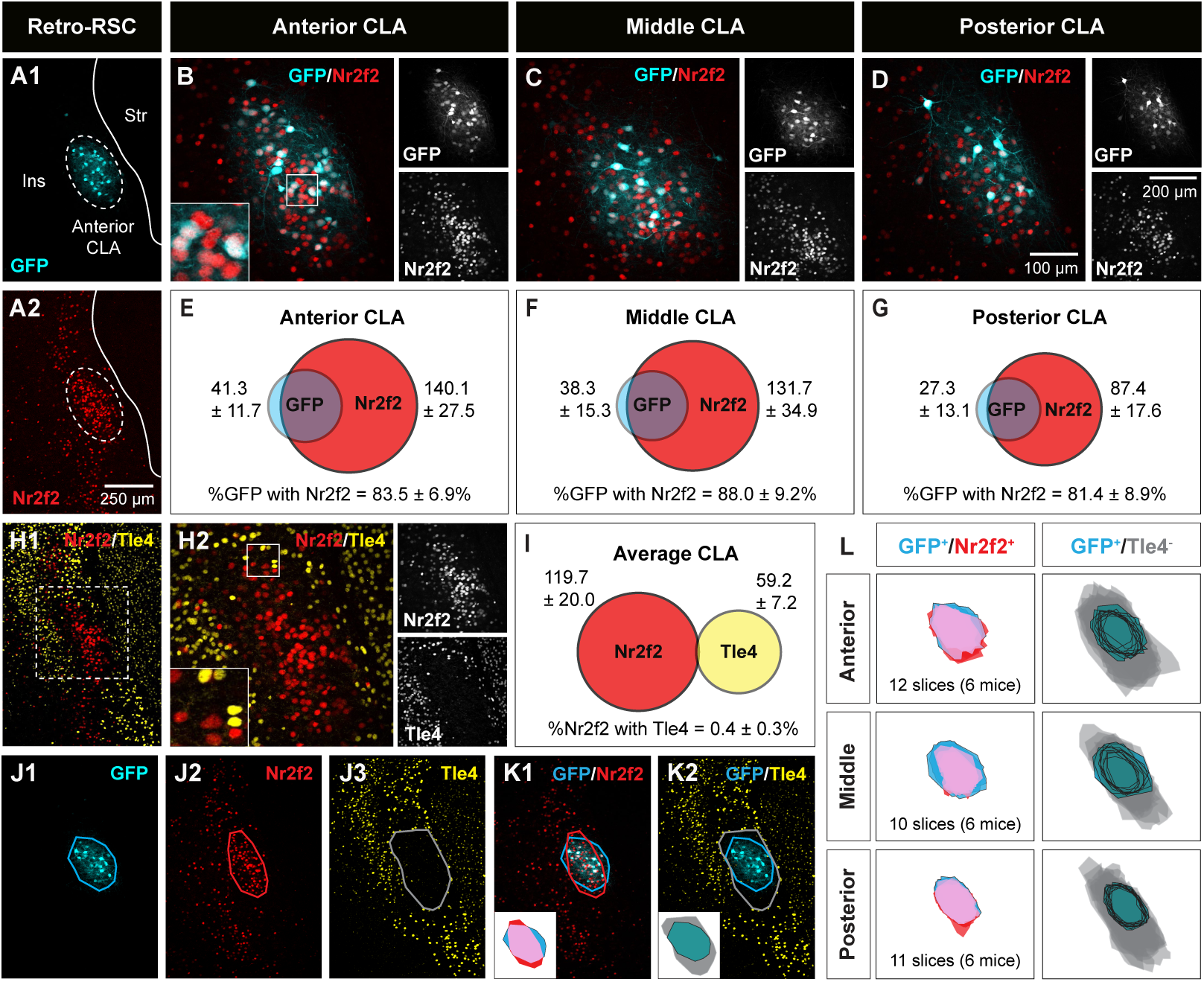
Spatial pattern of Nr2f2 and Tle4 expression enables localization of claustrum projection neurons. **A1-A2:** Representative images of the anterior claustrum showing retrograde GFP labeling **(A1)** and Nr2f2 expression **(A2)** following AAV injection into the retrosplenial cortex (Retro-RSC). Dashed ellipses highlight GFP expression in the claustrum (CLA) without labeling the surrounding insula cortex (Ins) or striatum (str). **B-D:** Representative images showing colocalization of GFP labeling with Nr2f2 in the anterior (**B**), middle (**C**) and posterior (**D**) CLA (left, merged imaging channels; right, single channel images). **E-G:** Venn diagrams representing the mean number of cells expressing Nr2f2 (red) and GFP (cyan), and colocalization of GFP with Nr2f2 in the anterior (**E**), middle (**F**), and posterior (**G**) CLA. Values indicate mean ± standard deviation from n = 6 mice (3 male and 3 female). **H1, H2:** Two representative images of the same coronal plane showing lack of colocalization between Nr2f2 and Tle4 in the CLA. **H2** is a magnified image of the dashed box in **H1**. For **H2**, merged imaging channels are on the left and single channel images are on the right. **I:** Venn diagram representing the mean number of cells expressing Nr2f2 (red) and Tle4 (yellow), and the colocalization of Nr2f2 with Tle4, averaged across anterior, middle and posterior CLA (n = 6 mice, 3 male and 3 female). **J1-J3:** Example images of the CLA region showing polygons outlining the boundaries of GFP cells and neuropil (cyan, **J1**), Nr2f2 concentric expression (red, **J2**), and absence of Tle4 expression (grey, **J3**). **K1-K2:** Overlay of polygons from **J1-J3.** Bottom-left insets show spatial registration of GFP outline with Nr2f2 outline (**K1**), and GFP outline with Tle4 outline (**K2**) (see Materials and Methods for details). Pink represents the topographic overlap of GFP/Nr2f2 expression, grey represents the region of low Tle4 expression. **L:** Cumulative spatial overlap of GFP^+^/Nr2f2^+^ labeling (left), and the spatial location of GFP labeling within the region of relative Tle4 absence (GFP^+^/Tle4^-^, right), in the anterior (top), middle (middle) and posterior (bottom) CLA across 6 mice. Insets in B, H2 (bottom-left) are 2.5× fold magnifications of areas in white boxes. Values in E, F, G, I represent mean ± standard deviation.

Outside of the claustrum, Nurr1 and Tle4 are found in deep layers of the sensory cortex, where Tle4 labels layers 5/6, and Nurr1 labels layer 6b (4,31–33). Therefore, we compared colocalization of Nurr1 with Tle4 in the claustrum to that in the cortex, namely in the motor and somatosensory cortices (**Figure 1P**). Markedly, while only a small number of Nurr1-labeled claustrum cells co-expressed Tle4 (∼1.5%) (**Figure 1Q, T, Supplementary Figure 2A, C, E**), the vast majority of Nurr1-labeled cells co-expressed Tle4 (>96%) in cortical layer 6b (**Figure 1R, S, U, V**). As such, this stark difference in Tle4 molecular profile between Nurr1-expressing cells in the claustrum-insula complex versus the cortex can be used to discriminate claustrum projection neurons from nearby neuronal populations in cortical layer 6b.

In addition to Nurr1, Nr2f2 is another cortical marker for claustrum cells (12,34). Similar to Nurr1, there was an extensive overlap between Nr2f2-immunoreactive cells and GFP-labeled RSC-projecting cells in the central zone (**Figure 2A1, A2**), and this overlap was maintained in claustrum subdivisions spanning the anteroposterior axis (>81%) (**Figure 2B-G**). However, GFP and Nurr1 co-expression was consistently higher than GFP and Nr2f2 co-expression across claustrum subdivisions (see **Figure 1**). There was very low co-expression of Nr2f2 and Tle4 in the claustrum region (∼0.4%) (**Figure 2H, I, Supplementary Figure 2B, D, F**), thus confirming lack of colocalization between Tle4 and claustrum projection neurons in the central zone.

In terms of spatial distribution, there was a dense group of cells in the claustrum central zone that exhibited strong Nr2f2 labelling when compared to the surrounding Nr2f2-immunoreactive cells, and this patch of Nr2f2-enriched cells seems to predominantly coincide with RSC-projecting claustrum cells (**Figure 2J1, J2, K1**). On the other hand, the ring-shaped clustering of Tle4-immunoreactive cells around RSC-projecting cells appears to demarcate the perimeter of the claustrum (**Figure 2J1, J3, K2**). The location of Nr2f2-enriched and Tle4-devoid domains relative to RSC-projecting cells was preserved across anterior, middle and posterior claustrum subdivisions, and was reproducible in different mice (**Figure 2L**). Accordingly, these findings suggest a cell-type-specific spatial organization in the claustrum, with strongly labeled Nr2f2-positive cells occupying a relatively central domain along the dorsoventral axis of the claustrum, i.e. the central zone, and Tle4-expressing cells defining the perimeter of the claustrum.

Collectively, our data indicates that that Tle4 enrichment is exclusive to structures abutting the claustrum. Our data also shows that Nurr1 and Nr2f2 label the majority of claustrum cells in the central zone, albeit their labelling profile fails to delineate the boundaries of the claustrum. As such, combining Nurr1/Nr2f2 with Tle4 labelling highlights the contrast in spatial patterning between Nurr1/Nr2f2 and Tle4 within the claustrum region, thus providing an exceedingly improved approximation of claustrum cell mapping.

### 4.2. Absence of Tle4 expression is a common feature for discrete subpopulations of claustrum cells

Are there neuronal populations devoid of Tle4 expression in the claustrum beside RSC-projecting neurons? To answer this question, we explored Tle4 colocalization with different subpopulations of claustrum projection neurons. It has been previously determined that claustrum neurons projecting to independent cortical regions are differentially distributed along the dorsoventral axis (15,16). To encompass the entire dorsoventral landscape of topographically positioned claustrum cells, we targeted different claustrocortical modules that are concentrated across the dorsoventral claustrum axis. To this end, we employed the same AAV approach as in the previous experiment to retrogradely trace claustrum cells projecting to the ACC (**Figure 3A, B**), MOp (**Figure 3G, H**) and LEC (**Figure 3M, N**), which comprise somewhat different claustrum populations in dorsal (**Figure 3C1**, **Figure 3I1**) and ventral zone (**Figure 3O1**) relative to the central zone we determined above (see **Figure 2**).

**Figure 3:**
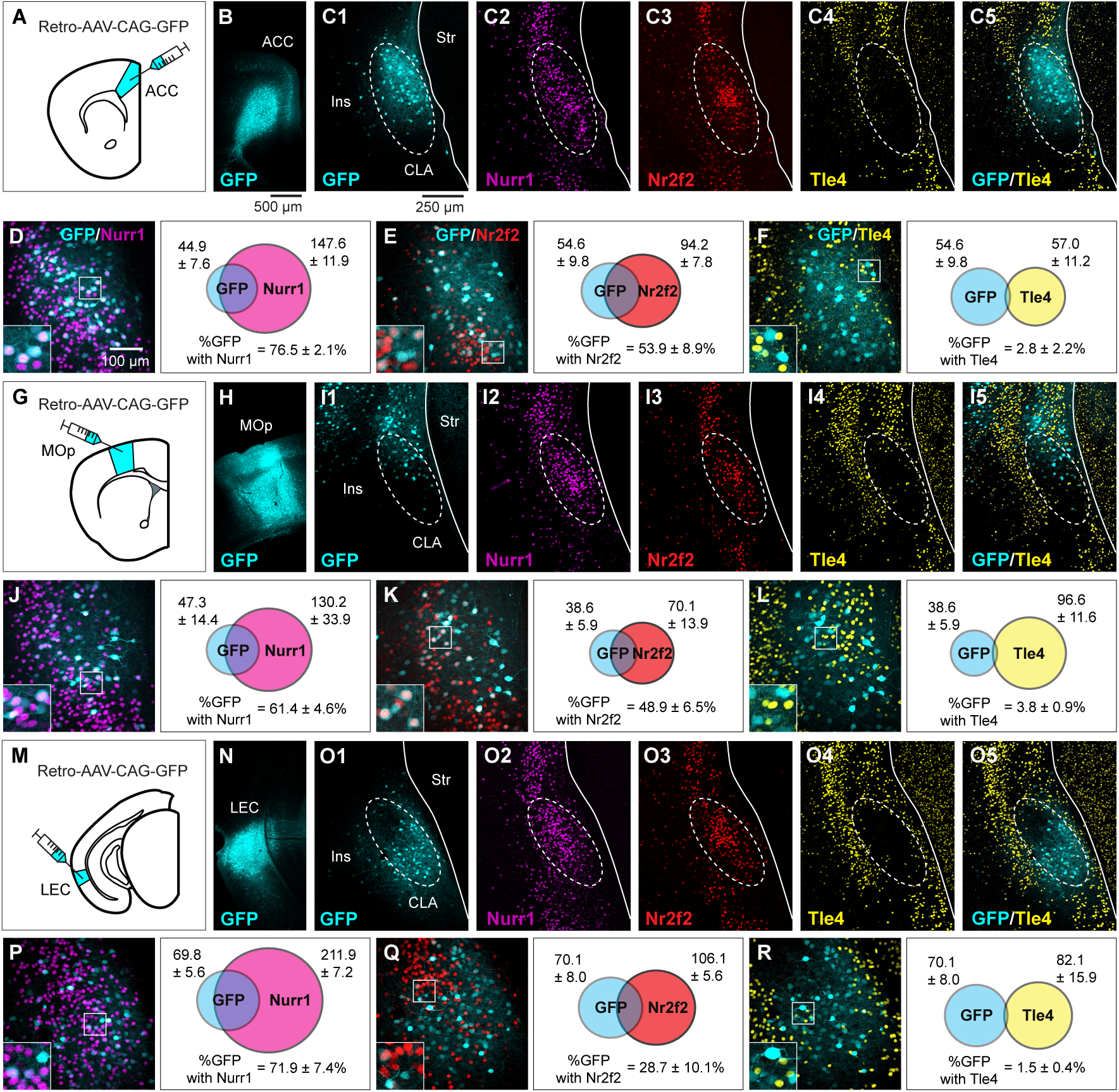
Claustrum neurons projecting to different cortical regions exhibit colocalization with Nurr1 and Nr2f2, but not with Tle4. **A:** Injection of retrograde AAV-CAG-GFP into the anterior cingulate cortex (ACC). Claustrum tissue was processed 14 days post injection. **B:** Example of an AAV injection site in the ACC. **C1-C4:** Representative images from coronal sections of the anterior claustrum (CLA) relative to the insula (Ins) and the striatum (Str) showing retrograde GFP labeling following AAV injection the ACC **(C1)**, along with the expression of Nurr1 **(C2)**, Nr2f2 **(C3),** Tle4 **(C4)**, and merged Tle4/GFP **(C5)**. All panels are from the same slice, except Nurr1 which is from an adjacent slice. Dashed ellipses highlight the region of low Tle4 expression. **D-F**: Representative images (left) and Venn diagrams (right) showing colocalization of GFP labeling with Nurr1 **(D)**, Nr2f2 **(E)** and Tle4 **(F)** in the anterior CLA. Panels E and F are the same slice. Venn diagrams in each panel show the mean number of cells expressing GFP with Nurr1 **(D)**, Nr2f2 **(E)**, and Tle4 **(F)** (n = 5 mice, all male). Values shown are the average cell counts across anterior, middle, and posterior planes of the CLA. Insets in left panels of D, E, F are 2.5× fold magnifications of areas in white boxes**. G-L** Same as **A-F** but for experiments with retrograde AAV-CAG-GFP injected into the primary motor cortex (MOp) (n = 4 mice, all male). **M-R:** Same as A-F, but for retrograde AAV-CAG-GFP injected into the lateral entorhinal cortex (LEC) (n = 4 mice, all male). Values in right panels of **D**, **E, F, J, K, L, P, Q, R** represent mean ± standard deviation.

We compared AAV labelling in each claustrocortical module with Nurr1, Nr2f2 and Tle4. For ACC-projecting cells, GFP expression was mainly confined between the central zone, demarcated by the enriched labelling of Nurr1 and Nr2f2 (**Figure 3C1-3**), and the dorsal zone, delineated by the upper limit of Tle4-devoid region (**Figure 3C4**, **C5**), thus corroborating our previous findings of ACC-projecting claustrum cells being preferentially located within central and dorsal zones (15). On average, ACC-projecting cells were enriched in Nurr1 (∼76%), and to a lower extent in Nr2f2 (∼54%) (**Figure 3D, E**), and they rarely expressed Tle4 (<3%) (**Figure 3F**). Correlating the position of MOp-projecting cells with Nurr1 and Nr2f2 dense patches showed that MOp module corresponds to dorsal parts of the claustrum (**Figure 3I1-I3**). Nevertheless, the Tle4 spatial distribution suggests that the module of MOp-projecting cells probably extends beyond the claustrum dorsal zone, raising the possibility that a subset of MOp-projecting cells is not part of the claustrum but rather belong to nearby dorsally located cortical structures (**Figure 3I4, I5**). Quantification for the MOp-projecting module indicated that on average ∼61% of the cells co-expressed Nurr1 (**Figure J3**), ∼54% of the cells co-expressed Nr2f2 (**Figure 3K**), and <4% of the cells co-expressed Tle4 (**Figure 3L**). These co-expression levels were somewhat similar to the ACC-projecting module.

LEC-projecting cells were clustered ventral to Nurr1 and Nr2f2 dense patches (**Figure 3O1-O3**), and yet, these cells were contained within the lower limit of the Tle4-devoid area (**Figure 3O4, O5**). This suggests that the LEC claustrocortical module is restricted to the ventral claustrum zone, which is in agreement with previously published data (15,35). While the colocalization of Nurr1 with LEC-projecting cells was comparable to ACC- and MOp-projecting cells (on average ∼72%, **Figure 3P**), Nr2f2 colocalized with LEC-projecting cells at a considerably low level relative to ACC- and MOp-projecting cells (on average ∼29%, **Figure 3Q**). Lastly, there was limited co-expression of LEC-projecting cells with Tle4 (on average ∼1.5%). Accordingly, Tle4 colocalization with all claustrocortical modules studied here was consistently low (<5%), suggestive of a prominent association between lack of Tle4 expression and claustrum projection neurons across diverse claustrocortical modules.

We took a closer look at colocalization of Nurr1, Nr2f2 and Tle4 with RSC, ACC, MOp and LEC claustrocortical modules in the anterior, middle and posterior claustrum subdivisions. Overall, cells in the RSC module exhibited the most robust Nurr1 and Nr2f2 expression among all modules, and this was consistent across all anteroposterior subdivisions (**Figure 4A-F, Supplementary Table 1**). In particular, differences between the RSC module and the ACC, MOp and LEC modules were significant in the anterior and middle claustrum with respect to Nurr1 (**Figure 4A, B**) and in all subdivisions with respect to Nr2f2 (**Figure 4D-F**). The only exception was Nurr1 expression in the posterior claustrum, where levels for the RSC module were significantly higher than in the MOp module, but not in ACC and LEC modules (**Figure 4C**). Conversely, Nurr1 expression in the MOp module was consistently lower than in the ACC and LEC modules across all anteroposterior subdivisions (**Figure 4A-C**). Yet, these differences were only significant in the anterior claustrum relative to ACC module (**Figure 4A**), and the posterior claustrum relative to ACC and LEC modules (**Figure 4C**). Also, Nr2f2 expression in the LEC module was lower than in the ACC and MOp modules (**Figure 4D-F, Supplementary Table 1**), with differences being significant only between LEC and ACC modules in the middle and posterior claustrum (**Figure 4E, F**). Thus, the expression of Nurr1 and Nr2f2 in claustrum cells varies between distinct claustrocortical modules, and also seems to be dependent on cell position along the anteroposterior axis. There were no significant differences in Tle4 levels between the different claustrocortical modules in any anteroposterior claustrum subdivision (**Figure 4G-I, Supplementary Table 1**). These results provide evidence that distinct cell subsets in the claustrum are overwhelmingly devoid of Tle4 expression regardless of their dorsoventral or anteroposterior position.

**Figure 4:**
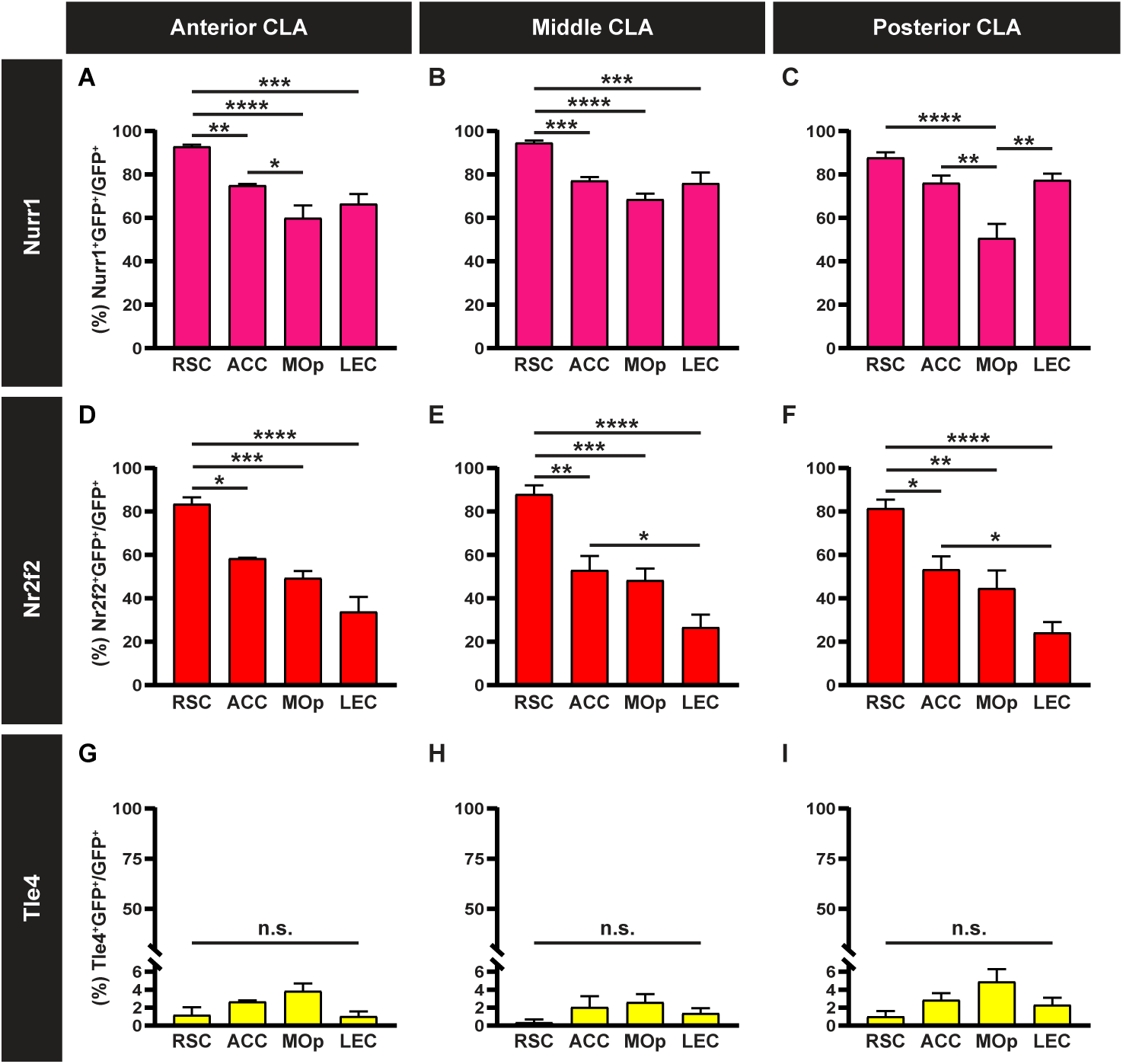
Claustrum neurons projecting to the retrosplenial cortex express Nurr1 and Nr2f2 more extensively than other claustrocortical pathways. **A-C:** Bar plots showing the mean percentage of GFP-expressing cells colocalized with Nurr1 (Nurr1^+^GFP^+^) in neurons projecting to the retrosplenial cortex (RSC), anterior cingulate cortex (ACC), primary motor cortex (MOp) and lateral entorhinal cortex (LEC) in the anterior **(A)**, middle **(B)** and posterior **(C)** claustrum (CLA). **D-F**: Same as A-C, but for GFP-expressing cells colocalized with Nr2f2 (Nr2f2^+^GFP^+^). **G-I:** Same as A-C, but for GFP-expressing cells colocalized with Tle4 (Tle4^+^GFP^+^). Error bars represent mean ± SEM; *p < 0.05, **p < 0.01, ***p < 0.001, ****p < 0.0001; one-way ANOVA followed by Bonferroni test (see **Supplementary Table 2** for detailed statistical analysis). RSC (n = 6 mice, 3 male and 3 female), ACC (n = 5 mice, all male), MOp (n = 4 mice, all male), LEC (n = 4 mice, all male).

In summary, by selectively tracing separate claustrocortical modules, we were able to determine that not all projection neurons express the claustrum-enriched markers Nurr1 and Nr2f2. Interestingly, this heterogeneity of Nurr1/Nr2f2 expression in the claustrum appears to be spatially specific, where cells projecting from the central zone to the RSC comprise the least heterogeneous cell population. Our experimental approach demonstrates an overall minimal overlap between claustrum cells and Tle4-expressing cells. Thus, we have effectively established Tle4 as a bona fide marker for segregating the claustrum from adjoining cortical areas.

### 4.3. Claustrum projection neurons lack the expression of inhibitory marker genes

It was previously reported that Nurr1-expressing cells in the claustrum are excitatory neurons (25). Because here we found that a sizeable portion of projection neurons in ACC, MOp and LEC claustrocortical modules did not express Nurr1 (30-40%), we sought to examine whether these cells have excitatory or inhibitory identity. For this purpose, we opted to focus on the ACC module since its spatial domain covers parts of the central and the dorsal zones (see **Figure 3**).

PV and somatostatin (SST) inhibitory neuronal subtypes are known to make up 50-60% of claustrum interneurons (15,36,37). Therefore, we performed co-immunolabelling of ACC-projecting cells in the claustrum with PV and SST markers. There was virtually no overlap between ACC-projecting cells and the expression of PV (∼0.04%; **Supplementary Figure 3A-D**) or of SST (∼0.05%; **Supplementary Figure 3E-H**). This suggests that projection neurons in the claustrum, identified using retrograde tracing, are predominantly excitatory neurons in line with many previous reports (24,27,38–40).

### 4.4. Tle4 molecular profile is suitable for identifying claustrum during early postnatal development

Conducting tracing experiments in juvenile rodents is considerably demanding. Therefore, using retrograde tracing to map claustrum projection neurons is less common during postnatal development than during adulthood. Previous studies reported that locating the claustrum via PV plexus labelling of claustrum core or distribution of myelinated fibers surrounding the claustrum are less reliable before the beginning of the third postnatal week (20). Based on Allen Institute mouse brain in situ hybridization database (29), Tle4 expression at early postnatal stages, i.e. P4, distinctly engulfs a small area in the ventrolateral cortex that is adjacent to the striatum, presumably the claustrum (**Supplementary Figure 1B**). Accordingly, we determined if the Nurr1/Tle4 molecular profile we used to identify claustrum cells in adult mice is suitable for locating the claustrum during neonatal development. To test this, we injected retro-AAV-CAG-TdTomato into the ACC of mice at P0, P7 or P14 and analyzed virus expression after 7 days at P7, P14 or P21, respectively (**Figure 5A, B1-B3**). It was logical to choose ACC as our injection site at such a young age since ACC retrograde tracing labels claustrum projection neurons at a higher density than RSC [our data and (20)]. In addition, it was recently shown that CLA projections to the ACC develop earlier than RSC projections (20). In order to compare the developmental results which used a seven-day virus expression time, an adult group was also injected at P49 and tissue was collected at P56 **(Figure 5A, B4)**. It is important to note that for P0, P7 and P14 mice, we adjusted injection volume to a smaller brain size to localize bolus delivery within the ACC and minimize non-specific viral spread to nearby regions (see Materials and Methods) (**Figure 5B1-B3**).

**Figure 5:**
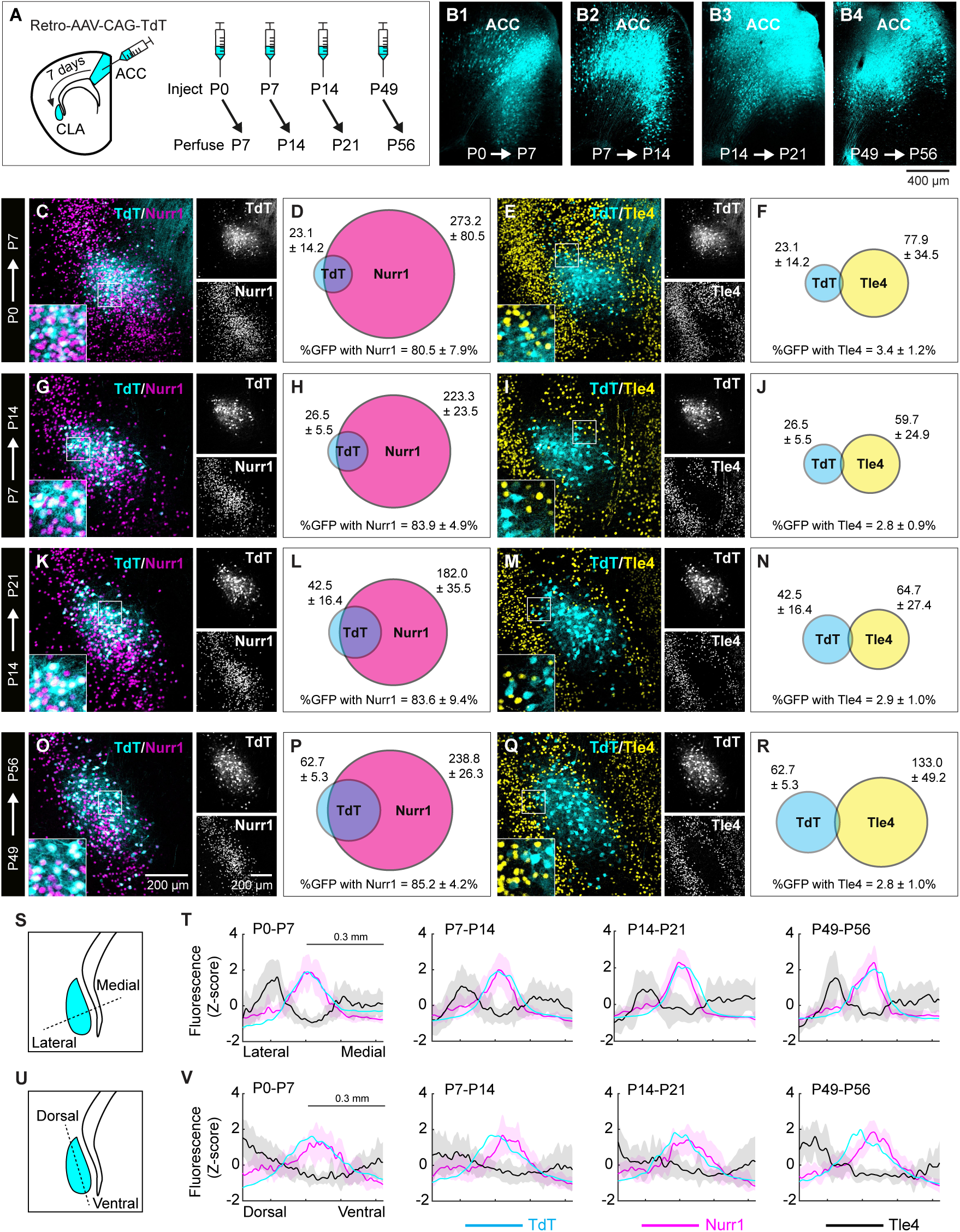
Claustrum cells in neonatal mice display the same Nurr1/Tle4 expression pattern as adult mice. **A**: Injection of retrograde AAV-CAG-TdT into the anterior cingulate cortex (ACC) at different postnatal days, followed by tissue collection 7 days post injection**. B1-B4:** Examples of AAV injection sites in the ACC at P7 **(B1)**, P14 **(B2)**, P21 **(B3)** and P56 **(B4)** following injection at P1, P7, P14 and P49, respectively**. C-F:** Representative images **(C, E)** showing the colocalization of TdTomato (TdT) labeling with Nurr1 **(C)** and Tle4 **(E)** in the anterior claustrum (CLA) at P7 (left, merged imaging channels; right, single channel images). Venn diagrams **(D, F)** representing the mean number of cells at P7 expressing Nurr1 and TdT **(D)**, or Tle4 and TdT **(F)** (n = 5 mice). Values shown are the average cell counts from the anterior, middle and posterior coronal planes of the CLA. **G-J:** Same as C-F, but for mice injected at P7 and perfused at P14 (n = 6 mice). **K-N**: Same as C-F but for mice injected at P14 and perfused at P21 (n = 7 mice). **O-R:** Same as C-F, but for mice injected at P49 and perfused at P56 (n = 5 mice). **S:** Quantification of fluorescence intensity in the CLA region (cyan) in the medial-lateral axis (dashed line). **T:** Measurement of fluorescence intensity (z-score) for TdT (cyan), Nurr1 (magenta) and Tle4 (black) labeling across the mediolateral axis of the CLA region at P7, P14, P21 and P56 (from left to right). Solid lines for each color represent the mean and light shaded areas of the same color represent standard deviation. **U-V**: Same as S-T, but for the dorsoventral axis. Insets in left panels of C, E, G, I, K, M, O, Q (bottom-left) are 2.5× fold magnifications of areas in white boxes. Values in D, F, H, J, L, N, P, R represent mean ± standard deviation (see **Supplementary Table 3** for details on mouse sex).

At P7-P21, the number of fluorescently labeled ACC-projecting cells was lower than at P56 in the claustrum region (on average ∼23 cells/slice at P7, ∼27 cells/slice at P14, ∼43 cells/slice at P21 and ∼63 cells/slice at P56), yet co-localization of TdTomato-tagged cells with Nurr1 and Tle4 at all developmental ages was comparable to P56: TdTomato-Nurr1 >80% at all ages; Tdtomato-Tle4 <5% at all ages (**Figure 5C-R**). Notably, the spatial distribution of most ACC-projecting cells at P7, P14 and P21 highly overlapped with Nurr1-enriched cell cluster in the claustrum region (**Figure 5C, G, K,**), and at the same time ACC-projecting cells occupied the Tle4-devoid zone that is surrounded by Tle4-expressing cells (**Figure 5E, I, M**). The same spatial patterns for both Nurr1 and Tle4 were replicated at P56 (**Figure 5O, Q**), and were identical to our results in adult mice using retro-AAV-CAG-GFP (see **Figure 3**). There was high colocalization of Nr2f2 and ACC-projecting cells between P7-P21 (>72%), which was on par with the colocalization level detected in P56 mice (>71%) (**Supplementary Figure 4**). These data suggest that AAC-projecting cells in the developing claustrum display an analogous molecular profile for Nurr1, Nr2f2 and Tle4 to their counterparts in the adult claustrum.

We further measured the spatial density for marker expression within the claustrum region at P7, P14 and P21 relative to P56. As expected, the peak density of ACC-projecting, i.e. TdTomato-expressing, cells largely coincided with Nurr1-expressing cells across the dorsoventral and mediolateral axes at all analyzed ages (**Figure 5S-V**). In contrast, the majority of Tle4-expressing cells were detected outside of the area with high TdTomato/Nurr1 fluorescence intensity from P7 and up to P56, with a bias towards being situated medial to as well as dorsal to the peak of TdTomato/Nurr1 fluorescence intensity (**Figure 5S-V**). Thus, at all early postnatal stages of claustrum development, the spatial density of Tle4 expression displayed an inverse trend to TdTomato and Nurr1 expression. These results demonstrate that spatial distribution of Nurr1 and Tle4 is nearly identical in neonatal (P7-P21) and adult mice (P56), suggestive of a uniform Nurr1/Tle4 expression pattern throughout postnatal claustrum development.

We compared Nurr1/Tle4 expression between the claustrum and cortical layer 6b during early postnatal development (**Supplementary Figure 5**). At P14, there was very low Nurr1/Tle4 colocalization in the claustrum (∼0.8%) (**Supplementary Figure 5B, E**), whereas layer 6b in the motor and somatosensory cortices displayed very high Nurr1/Tle4 colocalization (>97%) (**Supplementary Figure 5C, D, F, G**). These Nurr1/Tle4 colocalization results were very similar to what we found in adults (see **Figure 1**). Thus, claustrum and cortical layer 6b cells in the brain during postnatal development display the same Nurr1/Tle4 molecular profile as during adulthood. Altogether, similar to our findings in the adult claustrum, we were able to clearly identify claustrum projection neurons in neonatal mice by combining the expression profile of Nurr1-enriched cells with Tle4-devoid cells, thus rendering our approach a reliable choice for mapping claustrum location at different ages.

### 4.5. Selective recruitment of Nurr1-expressing cells over Tle4-expressing cells during CLA-mediated behaviours

Retrograde tracing of claustrum projection neurons allowed us to validate the molecular profile of Nurr1 and Tle4 in targeted claustrocortical modules. Therefore, we sought to test our Nurr1/Tle4 paradigm across widespread cell populations in the claustrum. Exposure to a novel context robustly activates claustrum neurons distributed across varied modules (27). To explore whether our combinatorial Nurr1/Tle4 approach could identify broadly located claustrum neurons, we placed adult mice in a novel OF, which has been previously shown to augment claustrum neuronal activity (26,27), and thereafter labeled activated claustrum neurons with the immediate early gene marker cFos (41,42). Baseline expression of cFos in littermates kept in their home cage, i.e. naive mice, served as our control.

In keeping with previously published work (26,27), cFos expression in the claustrum region was notably enhanced in the OF group relative to the naive group (p<0.0001) (**Figure 6A, F, K**). However, the spatial distribution of Nurr1-enriched (**Figure 6B, G**) and Tle4-devoid (**Figure 6C, H**) zones indicates that while the number of cFos-expressing cells was elevated in the claustrum following OF placement, cFos expression also increased in a large area that is somewhat ventrolateral to the claustrum (**Figure 6F-H**). This area presumably includes the insula, which suggests that the claustrum is activated together with nearby cortical structures upon exposure to novel circumstances. Investigating cFos co-expression with Nurr1 and Tle4 revealed two trends: First, cFos expression was increased in a substantial fraction of Nurr1-immunoreactive cells, where there was ∼15.6% upregulation of cFos in Nurr1-immunoreactive cells compared to controls (p<0.0001) (**Figure 6D, I, L**). On the other hand, there was a modest, albeit significant, rise in Tle4-immunoreactive cells co-expressing cFos as compared to controls (∼2.7% upregulation of cFos; p<0.01) (**Figure 6E, J, M**). This suggests that although being in a novel environment preferentially activates neurons within the claustrum, some nearby neurons outside of the claustrum are activated as well. Second, >60% of cFos-immunoreactive cells were also Nurr1-expressing cells (**Figure 6D, I, N**), whereas <6% of cFos-immunoreactive cells were also Tle4-expressing cells (**Figure 6E, J, O**), suggesting that in the claustrum region, novelty-induced neuronal activation is mainly exhibited by Nurr1-expressing cell rather than Tle4-expressing cells. Thus, in line with our hypothesis, characterization of Nurr1 and Tle4 immunoreactivity following behavioural activation of claustrum neurons helped disentangle activated neurons in the claustrum from adjacent areas in spite of the unanticipated change in neuronal activation outside of the claustrum. Overall, these findings, along with our previous data obtained by tracing disparate cortical-projecting claustrum cells, lead us to conclude that juxtaposing the expression of the claustrum-enriched gene Nurr1 with the cortical-enriched gene Tle4 is a reliable tool for identifying claustrum cells.

**Figure 6:**
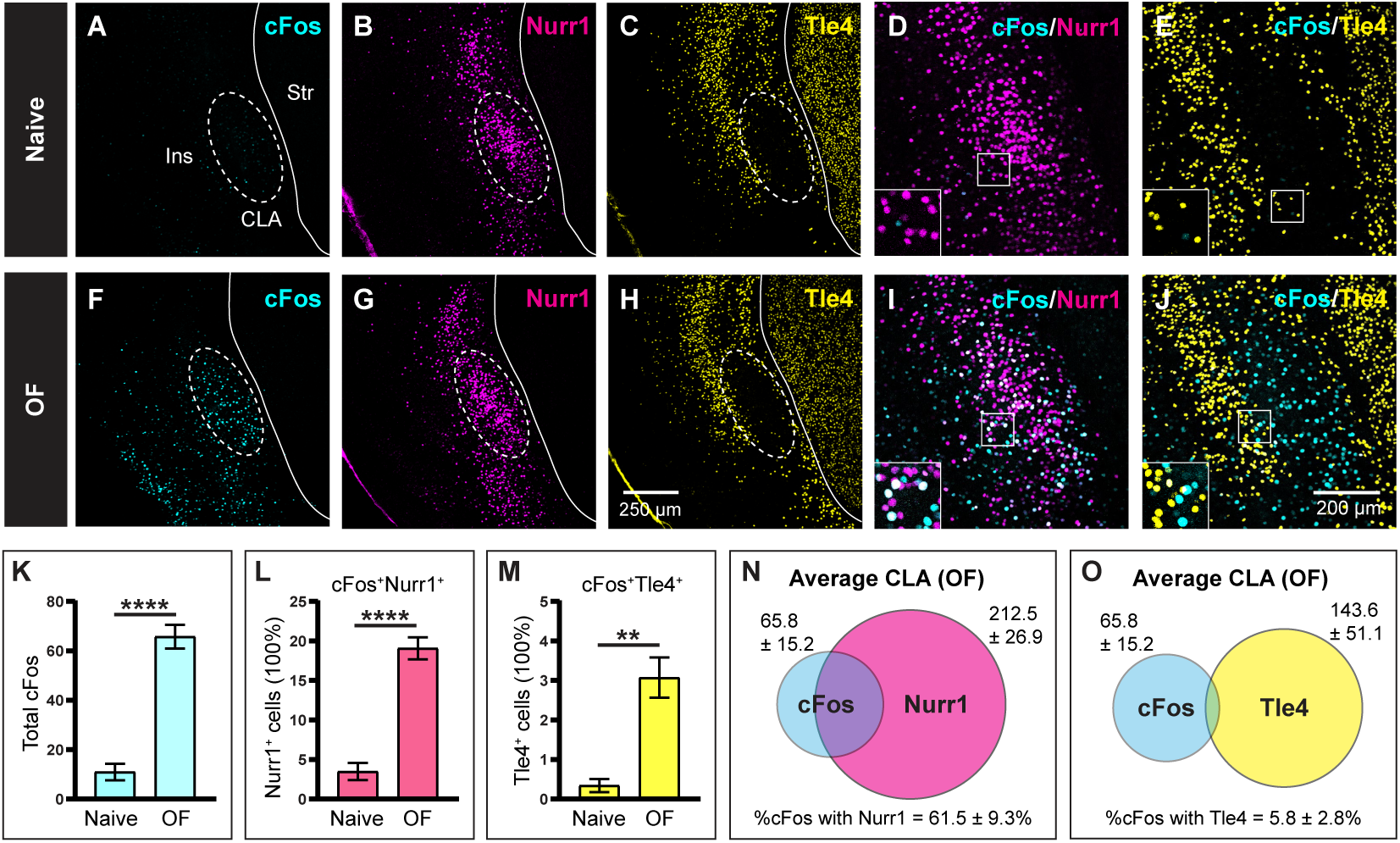
Exposure to a novel environment induces substantial activation of Nurr1-expressing cells and minimal activation of Tle4-expressing cells in the claustrum region. The marker for neuronal activation cFos was induced in the claustrum (CLA) by exposing mice to a novel open field (OF) for 10 minutes, followed by perfusion after 60-90 minutes. **A-C:** Representative images of the anterior CLA relative to the insula (Ins) and the striatum (Str) in naive control mice showing the expression of cFos **(A)**, Nurr1 **(B)** and Tle4 **(C)**. Dashed ellipses highlight absence of Tle4 expression in the CLA. **D-E:** Representative images showing colocalization of cFos with Nurr1 **(D)** and Tle4 **(E)** in the anterior CLA. **F-H:** Same as A-C but for mice exposed to OF. **I-J:** Same as D-E but for mice exposed to OF. **K-M:** Bar plots comparing naive and OF-exposed mice with respect to **(K)** the mean number of cFos-expressing cells, **(L)** the percentage of cFos-expressing cells colocalized with Nurr1 (cFos^+^Nurr1^+^) relative to the total number of Nurr1-expressing (Nurr1^+^) cells, and **(M)** the percentage of cFos-expressing cells colocalized with Tle4 (cFos^+^Tle4^+^) relative to the total number of Tle4-expressing (Tle4^+^) surrounding the CLA region. **N-O:** Venn diagrams representing the mean number of cells expressing cFos (cyan, **N, O**), Nurr1 (magenta, **N**) and Tle4 (yellow, **O**), along with the colocalization of cFos with Nurr1 **(N)**, and cFos with Tle4 **(O)** relative to the total number of cFos-expressing cells in the OF group. Values shown are the average cell counts across the anterior, middle and posterior planes of the CLA. Insets in D, E, I, J (bottom-left) are 2× fold magnifications of areas in white boxes. Error bars in K-M represent mean ± SEM; **p < 0.01, ****p < 0.0001; unpaired t-test: (K) t(13) = 7.53, p = 4.3×10^6^, (L): t(13) = 7.29, p = 6.1×10^6^, (M) t(13) = 3.68, p = 0.003. Values in N, O represent mean ± standard deviation. Naive group (n = 5 mice), OF group (n = 10 mice) (see **Supplementary Table 4** for details on mouse sex).

**Figure 7:**
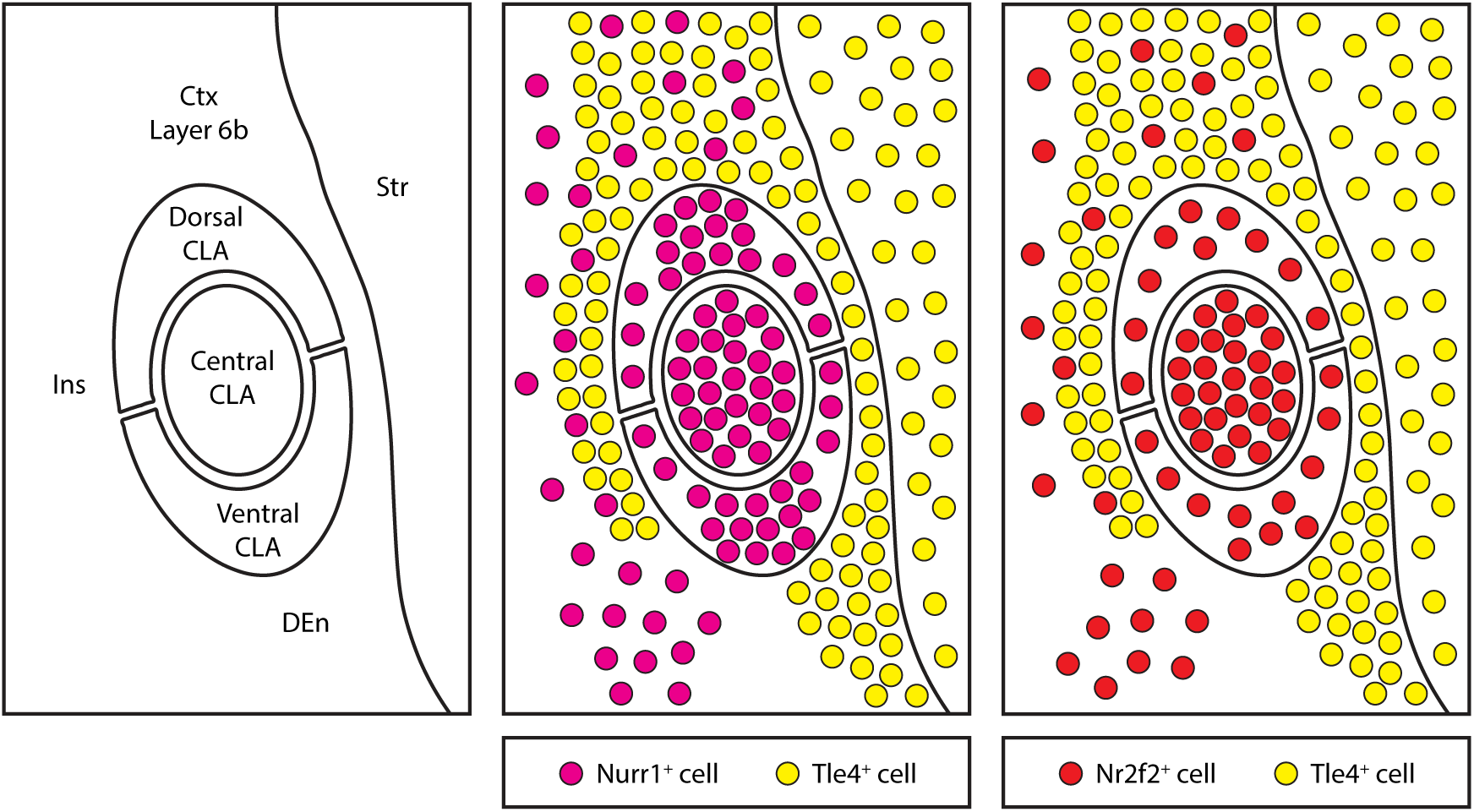
Schematic representation of mapping the claustrum using the claustrum-enriched markers Nurr1 and Nr2f2 in combination with the claustrum-devoid cortical marker Tle4. Nurr1-expressing (Nurr1^+^) and Nr2f2-expressing (Nr2f2^+^) cells are enriched in the claustrum, but they are also found in surrounding structures. While the dense patch of Nurr1^+^ cells is found in different claustrum zones, the dense patch of Nr2f2^+^ cells maps the central zone of the claustrum. Tle4-expressing (Tle4^+^) cells are devoid from the claustrum, and they are enriched in structures surrounding the claustrum. Juxtaposition of Nurr1^+^/Nr2f2^+^ cells with Tle4^+^ cells delineates claustrum borders. CLA: claustrum, Ctx: cortex, DEn: dorsal endopiriform nucleus, Ins: insula, Str: striatum.

## 5. DISCUSSION

In this study, we established a simplified, yet reliable, strategy to locate the mouse claustrum across development and adulthood. Our proposed method capitalizes on region-specific differences in the expression profile of typical claustrum markers versus cortical markers. The take-home message here is that the cortical marker Tle4 is ideal for this purpose. Indeed, spatial distribution of Tle4 expression distinctively avoids claustrum cells while strongly labelling cells that surround the claustrum (**Figure 7**). Other major findings of our work are the following: 1) Claustrum-enriched Nurr1 and Nr2f2 markers are not expressed equally in all claustrum cell populations, rendering these markers less useful for global identification of claustrum cells; 2) The topography of a strongly-labeled Nr2f2-psositive cell population can be used to map the central zone in the claustrum (**Figure 7**); and 3) The molecular profile of Nurr1 and Tle4 is maintained from early postnatal periods up to adulthood, and thus can be used to identify the claustrum during postnatal development.

Over the last two decades, a panel of claustrum markers were discovered, including genes like Netrin-G2, Latexin, Nr2f2 and Nurr1 (3,9,11–13). Although many of these markers are evolutionary conserved between primates and rodents, currently there is no claustrum marker that definitively outlines claustrum cells in rodents. One prominent claustrum marker is Nurr1, which has been instrumental for embryonic and postnatal developmental claustrum studies in rodents (10,11,13,20). Nevertheless, our retrograde labelling data indicate that despite Nurr1 being highly expressed in the claustrum central zone, a sizeable portion of cells in dorsal and ventral claustrocortical modules do not express Nurr1, and Nurr1 expression varies along the anteroposterior axis of the claustrum. Hence, Nurr1 should not be used as a “blanket” marker for all claustrum cells, but rather used as a potential marker for certain subtypes of claustrum neurons (9,43). Our results from cFos-labeled activated claustrum neurons corroborate this conclusion, where ∼38% of cFos-positive cells in the claustrum region lacked Nurr1 expression. We confirmed that claustrum projection neurons do not belong to either PV or SST subtypes of inhibitory neurons, suggesting that Nurr1-negative cells are likely to be another excitatory cell type or inhibitory NPY- and VIP-expressing cells (15,18,36).

Tle4 is a transcription factor mainly expressed by corticothalamic projection neurons in cortical layers 5 and 6 (32,33). A recent study determined that in the absence of Tle4 function during embryonic neurogenesis, cell progenitors that should give rise to corticothalamic projection neurons are misspecified, and instead adopt the identity of a different subtype of projection neurons (44). Throughout brain development, Nurr1 is highly expressed in the subplate, and later in adulthood, Nurr1 is found in cortical layer 6b, which is believed to be a remnant of the subplate (31,45). The claustrum has been speculated to share a common origin with the subplate/layer6b because they both co-express a number of genes, including Nurr1, and they have similar large-scale connectivity patterns (4). Given that Tle4 is required for the specification of corticothalamic projection neurons (44), the contrast in Tle4 expression we found between claustrum cells and Nurr1-expressing layer 6b cells suggests that the claustrum may have a different developmental trajectory than layer 6b (4). It would be interesting to examine the potential role of Tle4 in claustrum cell specification during embryogenesis.

In addition to showing that Tle4 expression is largely absent in claustrum cells, we reveal that the spatial distribution of Tle4 specifically encases the dorsal and lateral sides of the claustrum. This was evident in RSC, ACC and LEC retrograde labelling of claustrum projections. However, to our surprise, many MOp-projecting claustrum cells resided beyond the dorsal borders delineated by Tle4-immunoreactive cells. These MOp projections were outside of the concentric zone of Nurr1- and Nr2f2-labeled cells and showed lower levels of co-expression of Nurr1 and Nr2f2 than claustrum projections to other cortical regions. These results suggest that the MOp receives input from structures dorsally abutting the claustrum, potentially the gustatory and visceral cortices in addition to the claustrum. As for the claustrum boundaries in ventral areas, Tle4 does not clearly separate the claustrum from the embedded dorsal endopiriform nucleus. Yet, the dense patch of Nr2f2 expressing cells does not ventrally extend beyond the lower boundaries of the LEC clautrocortical module. Given that this module is biased towards the ventral zone of the claustrum (15), we speculate that spatial distribution of Nr2f2 avoids the dorsal endopiriform nucleus. Perhaps applying markers for the dorsal endopiriform nucleus, such as Ctgf, will confirm these observations (4,10,31). On the lateral side of the claustrum, our retrograde tracing and cFos-labelling data indicate that a thin sheet of Tle4-expressing cells located lateral to the claustrum represent a borderline that distinguishes claustrum projection neurons from laterally located structures, mainly the insula. The claustrum has been previously parcellated along the dorsoventral axis into claustrocortical modules that correspond to different populations of projection neurons (15). We provide evidence that the dense patch of Nr2f2-positive cells coincides with RSC-projecting neurons localized in the central zone (15,30). Thus, the expression profile of Nr2f2 could be used to identify the claustrum core, similar to PV neuropil and RSC-projecting claustrum cells. Combining Nr2f2 labelling together with Tle4 labelling can be utilized as a method for mapping dorsoventral claustrum zones, which could be an alternative to more invasive approaches that rely on retrograde tracing of claustrum projection neurons. Collectively, we determine that simultaneous labelling of Tle4 with Nurr1/Nr2f2 provides molecular landmarks that help locate claustrocortical neurons and separate them from other cells in the gustatory and visceral cortices on the dorsal side, from the dorsal endopiriform nucleus on the ventral side and from the insula on the medial side.

The postnatal development of the claustrum has been poorly studied to date. One reason for this is because mapping claustrum cells via retrograde tracing is challenging in mouse pups, and therefore tracing is seldom applied at early ages. By adapting our tracing protocol to newborn mice, we were able to employ the molecular profiles of Nurr1 and Tle4 to delineate the borders of the claustrum by the end of the first postnatal week. Other claustrum markers typically used in adulthood such as Oprk1, Ltx, Gnb4, and Snypr are not richly expressed until adolescence/adulthood (see **Supplementary Figure 1A**), which limit their use in a developmental context. In concordance with prior developmental studies (28), the relatively small brain size of P0/P1 mice necessitated lowering the injection volume. This consequently limited our retrograde tracing target selection to an area that receives claustrum projections in large numbers, provided that connections with the claustrum are established at an early postnatal age. A recent publication by Hoerder-Suabedissen et al. (20) showed that claustrum projections to the ACC reach peak density during the second postnatal week. Therefore, we chose ACC to be our target injection site in newborn mice. However, in contrast to Hoerder-Suabedissen et al. (20), we found that by the end of the third postnatal week, the number of ACC-projecting claustrum cells was much lower than in young adults. This could be attributed to the smaller injection volumes we used here. Still, our approach for neonates revealed that the expression pattern and spatial distribution of Nurr1 and Tle4 are highly selective to the claustrum, and these results were nearly identical to adults. Although Hoerder-Suabedissen et al. (20) identified different strategies for studying claustrum postnatal development, including PV-labelling plexus and differences in regional myelination, none of these strategies were suitable for locating the claustrum before the beginning of the third postnatal week. Therefore, the combinatorial Nurr1/Tle4 method we developed here has the advantage of mapping claustrum cells during the first two postnatal weeks.

Based on molecular labelling of neuronal activation via the intermediate-early gene cFos, it is assumed that claustrum cell activity increases during exposure to novelty (26,27). Consistently, our work indicates that being in a novel environment induces cFos expression at high levels in the Nurr1-enriched/Tle4-devoid region, and particularly in Nurr1-positive cells. These observations suggest that claustrum cells are indeed recruited upon encountering a novel context. Nevertheless, the small elevation of cFos labelling in Tle4-positive cells, and the detection of cFos expression lateral to Tle4-positive cells bordering the claustrum, indicate that nearby cortical structures, including the insula, are also engaged during novelty exposure. Accordingly, our Nurr1/Tle4 combinatorial approach is effective at distinguishing claustrum cells from their surroundings during claustrum-dependent behaviours, thus it is well suited for functional studies of the claustrum.

The small size of the claustrum, its hidden location between the much bigger neighboring structures striatum and insula, and the apparent anatomical continuity between claustrum cells and surrounding cortical cells have hampered our understanding of claustrum functions. Our dual-labelling approach of claustrum markers combined with cortical markers provides insights into previously unrecognized anatomical features of the claustrum. This approach may be a useful alternative for studying the anatomical and functional properties of the claustrum across different ages. Our work therefore paves the way for future studies to discern claustrum-specific anatomical and functional properties, whether during early postnatal development or adulthood.

## 6. LIST OF ABBREVIATIONS

ACC: Anterior cingulate cortex
AAV: Adeno-associated virus
CLA: Claustrum
Ctx: Cortex
DEn: dorsal endopiriform nucleus
Ins: Insula
L6b: Layer 6b
LEC: Lateral entorhinal cortex
MOp: Primary motor cortex
OF: Open field
P: Postnatal day
PBS: Phosphate buffer solution
PFA: Paraformaldehyde
PV: Parvalbumin
RSC: Retrosplenial cortex
SST: Somatostatin
Str: Striatum
TdT: TdTomato.

## 7. SUPPLEMENTARY INFORMATION

**Supplementary Figure 1:**
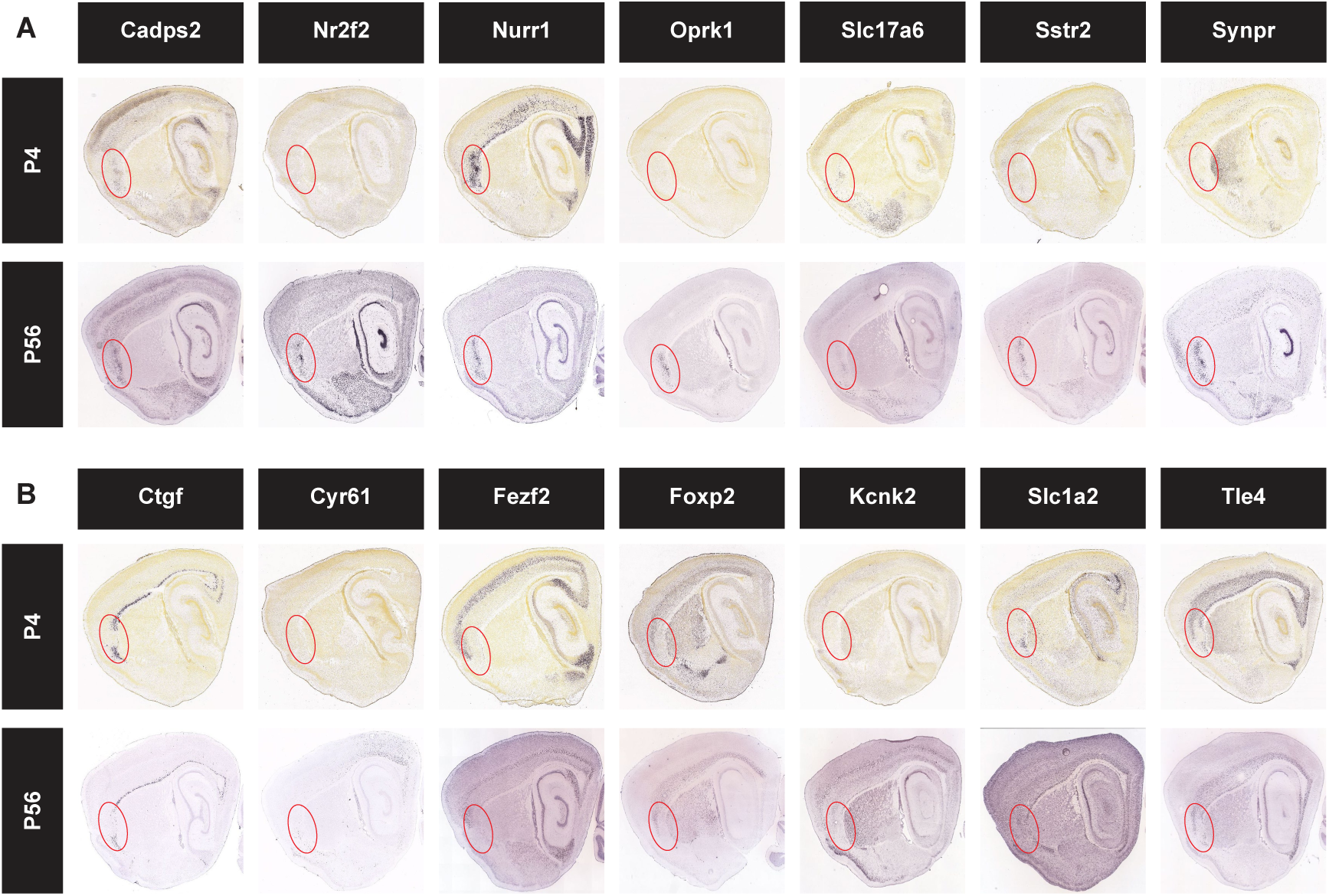
Selection of candidate marker genes that are absent in claustrum cells and at the same time enriched in other nearby cells. **A-B:** Chromogenic *in situ* hybridization images at P4 (top) and P56 (bottom) of claustrum-enriched genes **(A)** and cortical-enriched genes **(B)** in the claustrum region (red ellipse). Candidate genes were selected based on earlier transcriptomic data showing differential expression of these genes in claustrum cells versus cortical cells (23). Images were obtained from Allen mouse brain in situ hybridization database (http://developingmouse.brain-map.org).

**Supplementary Figure 2:**
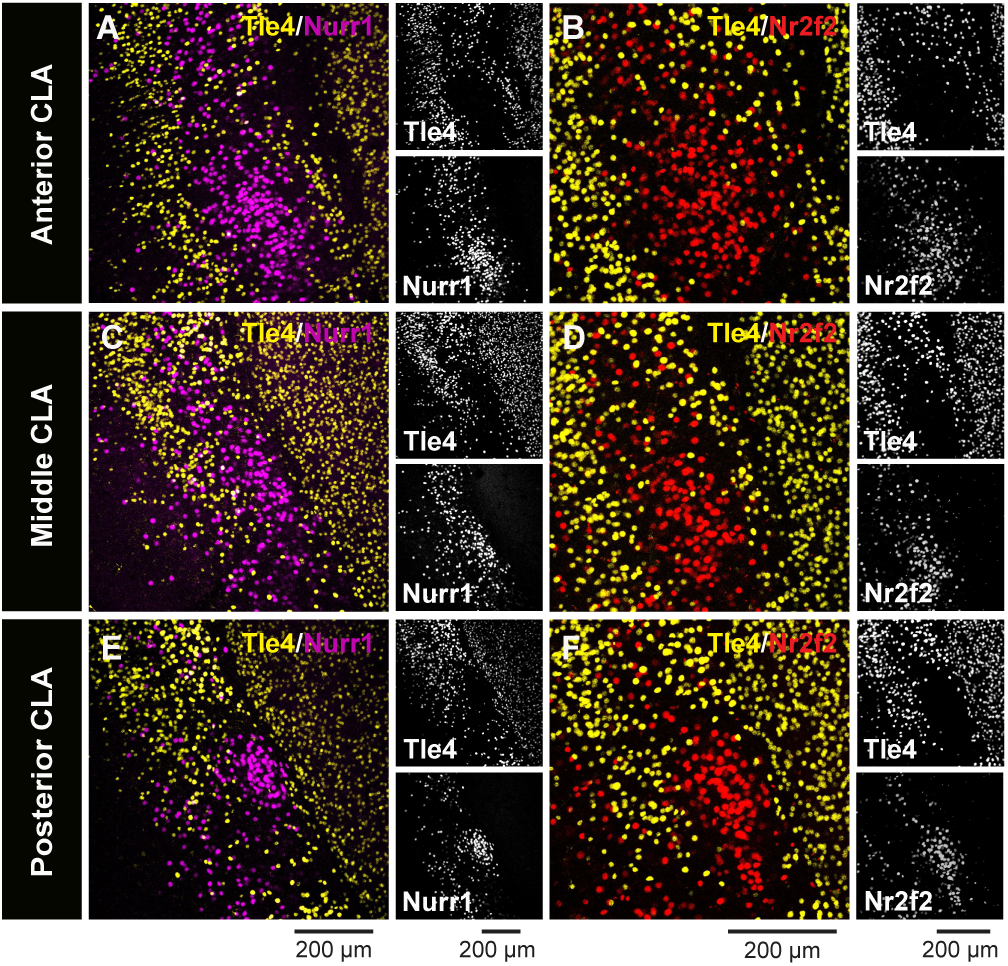
Expression of the claustrum-enriched markers Nurr1 and Nr2f2 relative to the expression of Tle4 across the anteroposterior axis of the claustrum. **A-F:** Representative images of the anterior **(A, B)**, middle **(C, D)** and posterior **(E, F)** claustrum showing co-labeling of Nurr1 **(A, C, E)** and Nr2f2 **(B, D, F)** with Tle4 (left, merged imaging channels; right, single channel images).

**Supplementary Figure 3:**
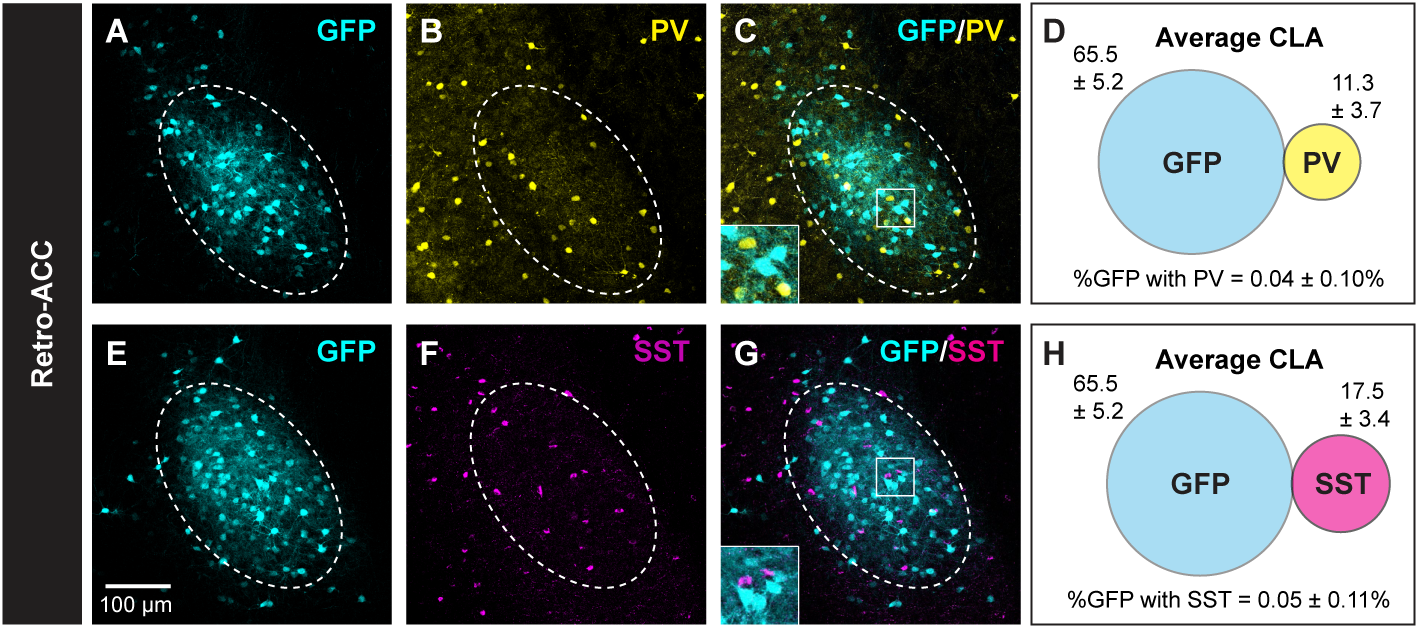
Lack of colocalization between claustrum projection neurons and inhibitory markers. **A-C:** Representative images of the anterior claustrum (CLA, dashed ellipses) showing retrograde GFP labeling **(A**), PV expression **(B)**, merged A and B channels **(C)** following AAV injection into the anterior cingulate cortex (Retro-ACC). **D:** Venn diagram representing the mean number of cells expressing GFP (cyan) and PV (yellow), along with the colocalization of GFP with PV relative to the total number of GFP-expressing cells. Values shown are the average cell counts from the anterior, middle and posterior planes of the CLA. **E-H:** Same as A-D, respectively, but for SST (magenta). Insets in C, G (bottom-left) are 2× fold magnifications of areas in white boxes. Values in D, H represent mean ± standard deviation (n = 5 mice, all male).

**Supplementary Figure 4:**
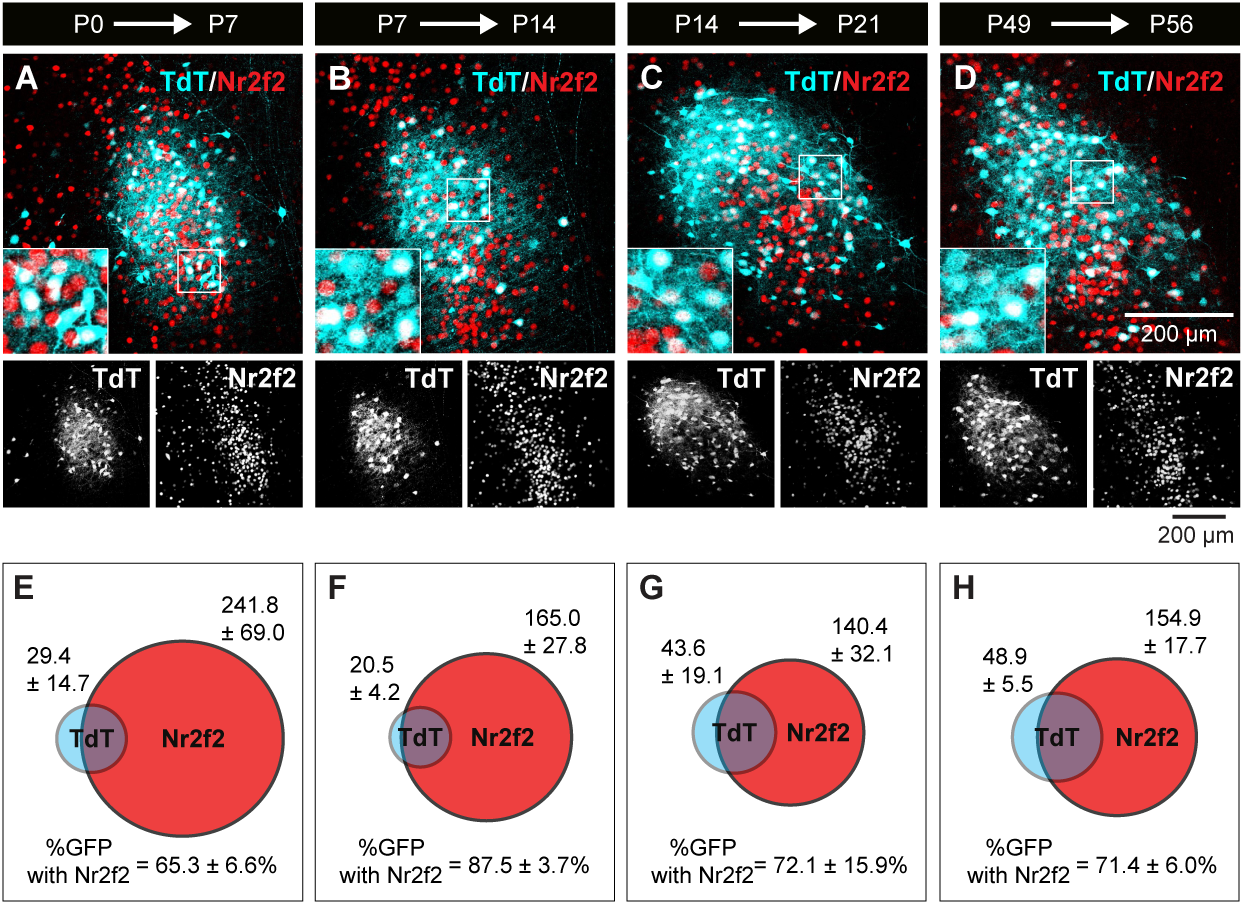
Nr2f2 displays high colocalization with ACC-projecting claustrum cells consistently throughout postnatal development. **(A-H)** Same experimental design as in Figure 5. **A-D**: Representative images showing colocalization of TdTomato (TdT) labeling with Nr2f2 in the anterior claustrum at P7 **(A)**, P14 **(B)**, P21 **(C)** and P56 **(D)** (left, merged imaging channels; right, single channel images). Insets (bottom-left) are 2.5× fold magnifications of areas in white boxes. **E-H:** Venn diagrams representing the mean number of cells expressing TdT (cyan) and Nr2f2 (red), along with the colocalization of TdT with Nr2f2 relative to the total number of TdT-expressing cells, at P7 **(E)**, at P14 **(F)**, at P21 **(G)** and at P56 **(H)**. Values shown represent mean ± standard deviation, which were calculated by averaging cell counts from the anterior, middle and posterior planes of the CLA (P7: n = 5 mice, P14: n = 4 mice, P21: n = 6 mice, P56: n = 3 mice) (see **Supplementary Table 3** for details on mouse sex).

**Supplementary Figure 5:**
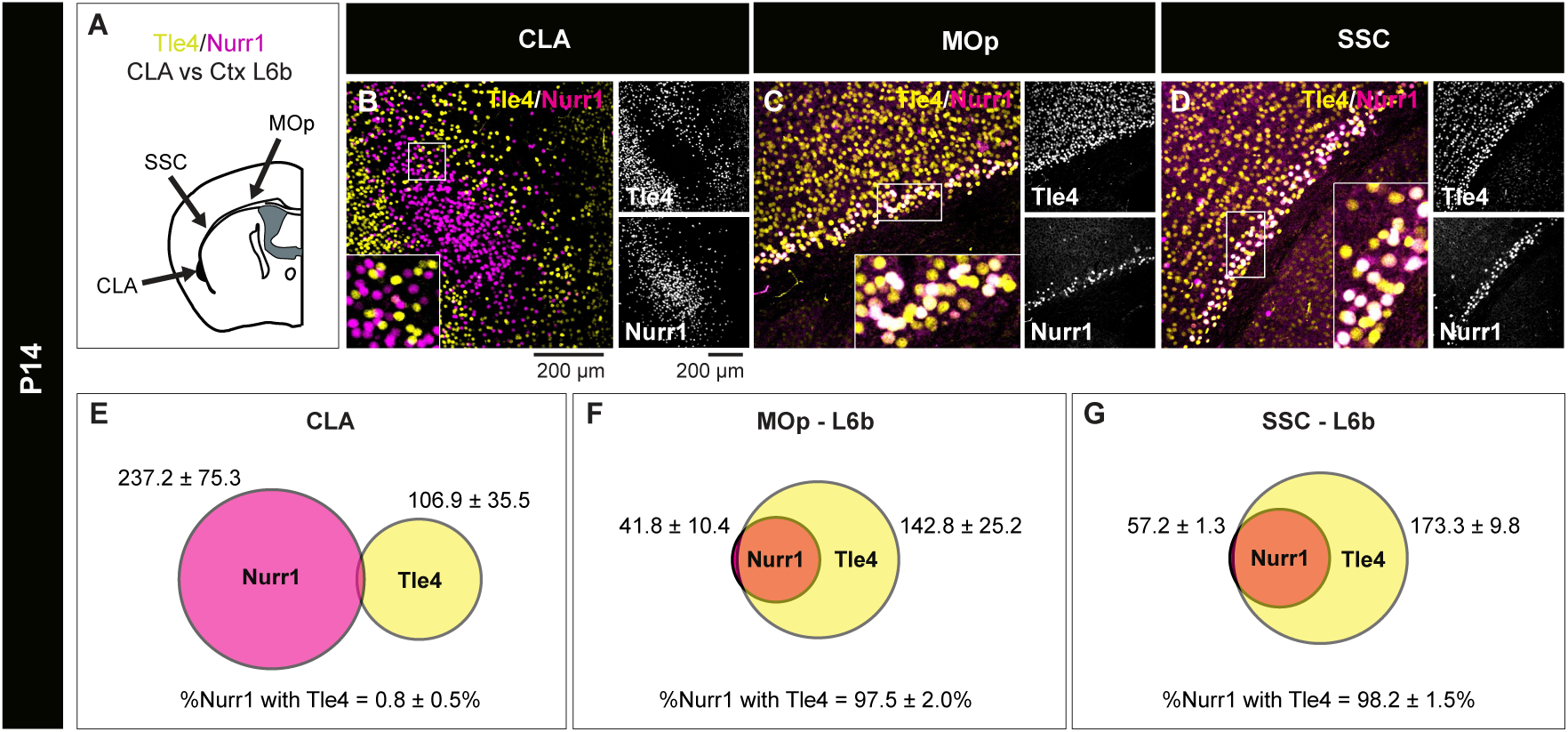
Colocalization of Nurr1 with Tle4 in the claustrum is different from that in the cortex at P14. **A:** Schematic of the experimental design at P14 comparing Nurr1 and Tle4 colocalization in the claustrum (CLA) to cortical layer 6b (Ctx L6b) in the primary motor cortex (MOp) and the somatosensory cortex (SSC). **B-D:** Representative images showing Nurr1 and Tle4 expression in the CLA **(B)**, the MOp **(C)** and the SSC **(D)** (left, merged imaging channels; right, single channel images). Insets are 2.5× fold magnifications of areas in white boxes. **E-G:** Venn diagrams representing the mean number of cells expressing Nurr1 (magenta) and Tle4 (yellow), along with the colocalization of Nurr1 with Tle4 relative to the total number of Nurr1-expressing cells in the CLA **(E)** (n = 6 mice, 3 male and 3 female), and Ctx L6b in the MOp **(F)** and in the SSC **(G)** (n = 3 mice, 2 male and 1 female) (See **Supplementary Table 1** for details on mouse sex). Insets in B, C, D are 2.5× fold magnifications of areas in white boxes. Values shown represent mean ± standard deviation. The mean for the CLA was calculated by averaging cell counts from the anterior, middle and posterior planes of the CLA region. The mean for Ctx L6b in MOp and SSC was calculated by averaging cell counts from slices on the same plane as the anterior, middle and posterior regions of the CLA.

**Supplementary Table 1:** Individualized marker quantification and colocalization in the claustrum (CLA), primary motor cortex (MOp), and somatosensory cortex (SSC) for each mouse at different postnatal ages. Values represent mean cell count per slice. For the claustrum, the mean was calculated by averaging data across the anterior, middle, and posterior subdivisions. The first column reflects mouse age, where P(x) indicates postnatal day when tissue was collected. For each age, mice were derived from ≥ 2 litters. F: female, M: Male.

| Mouse ID | Sex | Region | Nurr1+ | Tle4+ | Nurr1+/Tle4+ |
| --- | --- | --- | --- | --- | --- |
| P14-1 | F | CLA | 328.6 | 113.8 | 5.0 |
|  |  | MOp | 38.5 | 159.8 | 38.3 |
|  |  | SSC | 56.0 | 164.8 | 54.5 |
| P14-2 | F | CLA | 258.7 | 118.2 | 3.0 |
|  |  | MOp | N/A | N/A | N/A |
|  |  | SSC | N/A | N/A | N/A |
| P14-3 | F | CLA | 110.5 | 58.0 | 0.5 |
|  |  | MOp | N/A | N/A | N/A |
|  |  | SSC | N/A | N/A | N/A |
| P14-5 | M | CLA | 288.0 | 163.3 | 2.8 |
|  |  | MOp | 33.5 | 113.8 | 32.8 |
|  |  | SSC | 58.5 | 184.0 | 58.5 |
| P14-6 | M | CLA | 230.0 | 104.7 | 0.5 |
|  |  | MOp | 53.5 | 154.8 | 51.0 |
|  |  | SSC | 57.0 | 171.3 | 55.5 |
| P14-7 | M | CLA | 207.7 | 83.7 | 0.7 |
|  |  | MOp | N/A | N/A | N/A |
|  |  | SSC | N/A | N/A | N/A |
| P75-1 | F | CLA | 163.4 | 69.6 | 0.6 |
|  |  | MOp | 42.3 | 133.8 | 41.8 |
|  |  | SSC | 69.0 | 158.5 | 68.3 |
| P75-2 | F | CLA | 177.8 | 98.0 | 0.6 |
|  |  | MOp | 54.3 | 175.5 | 54.3 |
|  |  | SSC | 77.8 | 191.3 | 75.5 |
| P75-3 | M | CLA | 210.3 | 78.0 | 2.7 |
|  |  | MOp | 32.5 | 106.3 | 32.5 |
|  |  | SSC | 57.5 | 134.5 | 56.3 |
| P75-4 | M | CLA | 174.0 | 88.0 | 5.3 |
|  |  | MOp | 17.8 | 98.3 | 17.8 |
|  |  | SSC | 36.3 | 101.0 | 33.5 |

**Supplementary Table 2:**
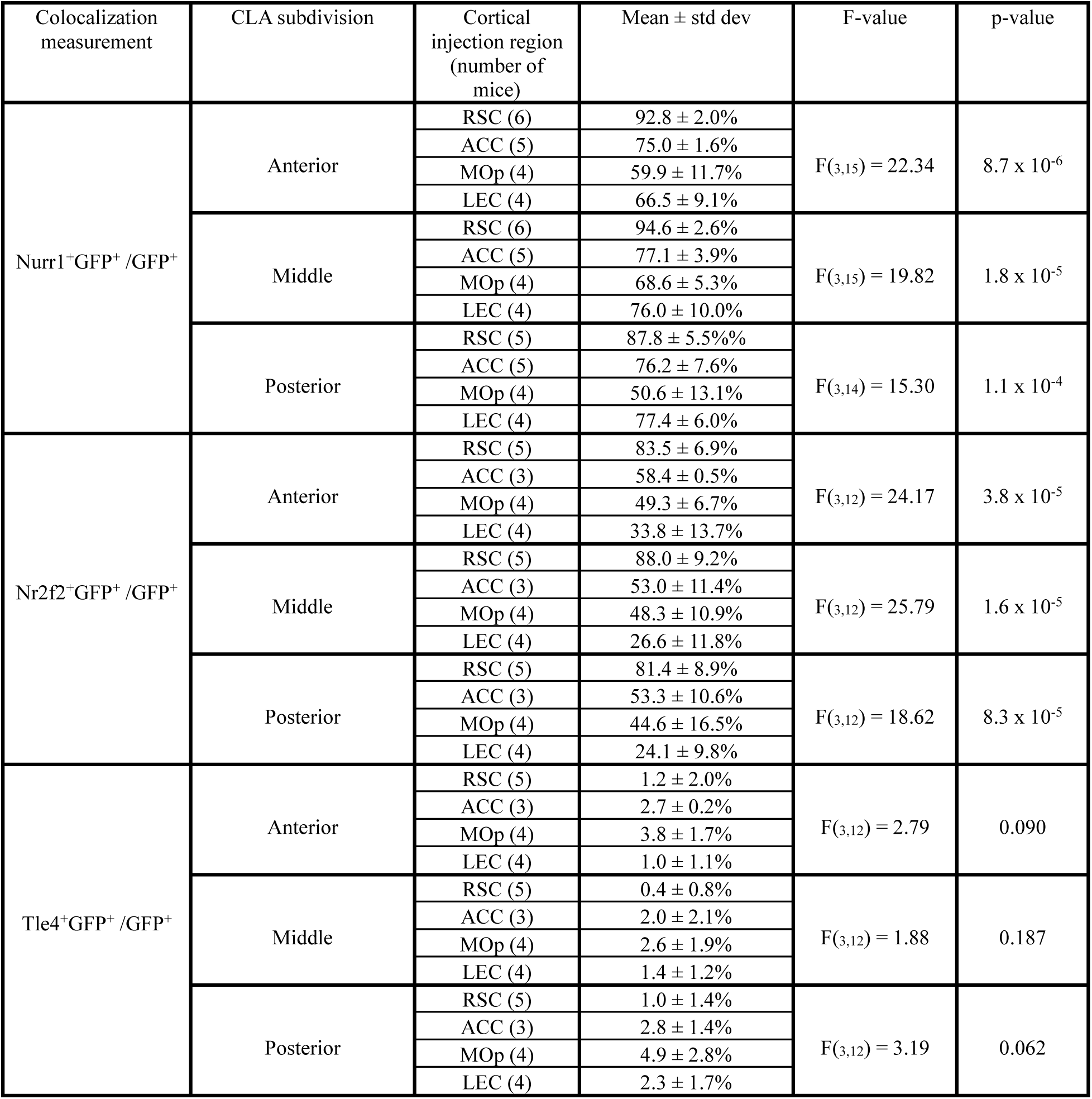
Quantification and one-way ANOVA analysis of marker colocalization with GFP^+^ cells in different claustrocortical anteroposterior subdivisions of the claustrum (CLA), following retrograde tracing from different cortical regions. ACC: anterior cingulate cortex, MOp: primary motor cortex, LEC: lateral entorhinal cortex, RSC: retrosplenial cortex, std dev: standard deviation. Mean ± std dev values represent the percentage of marker colocalization with GFP^+^ cells relative to total GFP^+^ cells.

| Colocalization measurement | CLA subdivision | Cortical injection region (number of mice) | Mean ± std dev | F-value | p-value |
| --- | --- | --- | --- | --- | --- |
| Nurr1<sup>+</sup>GFP<sup>+</sup> /GFP<sup>+</sup> | Anterior | RSC (6) | 92.8 ± 2.0% | F<sub>(3,15)</sub> = 22.34 | 8.7 x 10<sup>-6</sup> |
| | | ACC (5) | 75.0 ± 1.6% | | |
| | | MOp (4) | 59.9 ± 11.7% | | |
| | | LEC (4) | 66.5 ± 9.1% | | |
| | Middle | RSC (6) | 94.6 ± 2.6% | F<sub>(3,15)</sub> = 19.82 | 1.8 x 10<sup>-5</sup> |
| | | ACC (5) | 77.1 ± 3.9% | | |
| | | MOp (4) | 68.6 ± 5.3% | | |
| | | LEC (4) | 76.0 ± 10.0% | | |
| | Posterior | RSC (5) | 87.8 ± 5.5% | F<sub>(3,14)</sub> = 15.30 | 1.1 x 10<sup>-4</sup> |
| | | ACC (5) | 76.2 ± 7.6% | | |
| | | MOp (4) | 50.6 ± 13.1% | | |
| | | LEC (4) | 77.4 ± 6.0% | | |
| Nr2f2<sup>+</sup>GFP<sup>+</sup> /GFP<sup>+</sup> | Anterior | RSC (5) | 83.5 ± 6.9% | F<sub>(3,12)</sub> = 24.17 | 3.8 x 10<sup>-5</sup> |
| | | ACC (3) | 58.4 ± 0.5% | | |
| | | MOp (4) | 49.3 ± 6.7% | | |
| | | LEC (4) | 33.8 ± 13.7% | | |
| | Middle | RSC (5) | 88.0 ± 9.2% | F<sub>(3,12)</sub> = 25.79 | 1.6 x 10<sup>-5</sup> |
| | | ACC (3) | 53.0 ± 11.4% | | |
| | | MOp (4) | 48.3 ± 10.9% | | |
| | | LEC (4) | 26.6 ± 11.8% | | |
| | Posterior | RSC (5) | 81.4 ± 8.9% | F<sub>(3,12)</sub> = 18.62 | 8.3 x 10<sup>-5</sup> |
| | | ACC (3) | 53.3 ± 10.6% | | |
| | | MOp (4) | 44.6 ± 16.5% | | |
| | | LEC (4) | 24.1 ± 9.8% | | |
| Tle4<sup>+</sup>GFP<sup>+</sup> /GFP<sup>+</sup> | Anterior | RSC (5) | 1.2 ± 2.0% | F<sub>(3,12)</sub> = 2.79 | 0.090 |
| | | ACC (3) | 2.7 ± 0.2% | | |
| | | MOp (4) | 3.8 ± 1.7% | | |
| | | LEC (4) | 1.0 ± 1.1% | | |
| | Middle | RSC (5) | 0.4 ± 0.8% | F<sub>(3,12)</sub> = 1.88 | 0.187 |
| | | ACC (3) | 2.0 ± 2.1% | | |
| | | MOp (4) | 2.6 ± 1.9% | | |
| | | LEC (4) | 1.4 ± 1.2% | | |
| | Posterior | RSC (5) | 1.0 ± 1.4% | F<sub>(3,12)</sub> = 3.19 | 0.062 |
| | | ACC (3) | 2.8 ± 1.4% | | |
| | | MOp (4) | 4.9 ± 2.8% | | |
| | | LEC (4) | 2.3 ± 1.7% | | |

**Supplementary Table 3:** Individualized marker quantification and colocalization with claustrum cells that express Tdtomato (TdT) for each mouse at different postnatal stages. AAVretro-CAG-TdT was injected into the anterior cingulate. The values represent mean cell count within coronal slices, averaged across the anterior, middle, and posterior claustrum for individual mice. The first column reflects mouse age, where P(x) indicates postnatal day when tissue was collected. For each age, mice were derived from ≥ 2 litters. F: female, M: Male. Datasets 1 and 2 were either derived from adjacent slices or different mice.

| Mouse ID | Sex | TdT+<br>(dataset 1) | Nurr1+<br>(dataset 1) | TdT+<br>/Nurr1+<br>(dataset 1) | Tle4+<br>(dataset 1) | TdT+/Tle4+<br>(dataset 1) | TdT+<br>(dataset 2) | Nr2f2+<br>(dataset 2) | TdT+/Nr2f2+<br>(dataset 2) |
| --- | --- | --- | --- | --- | --- | --- | --- | --- | --- |
| P7-1 | F | 40.8 | 320.0 | 27.6 | 125.0 | 1.6 | 41.0 | 290.2 | 23.2 |
| P7-2 | F | 36.3 | 309.2 | 30.2 | 96.0 | 1.5 | N/A | N/A | N/A |
| P7-3 | M | 13.4 | 213.0 | 10.6 | 47.0 | 0.2 | N/A | N/A | N/A |
| P7-4 | M | 11.5 | 165.5 | 9.8 | 42.3 | 0.5 | N/A | N/A | N/A |
| P7-5 | M | 13.4 | 358.2 | 11.8 | 79.0 | 0.4 | 13.5 | 149.5 | 8.8 |
| P7-6 | M | N/A | N/A | N/A | N/A | N/A | 15.0 | 204.5 | 11.3 |
| P7-8 | M | N/A | N/A | N/A | N/A | N/A | 31.6 | 324.2 | 20.2 |
| P7-9 | M | N/A | N/A | N/A | N/A | N/A | 45.8 | 240.4 | 30.4 |
| P14-1 | F | 30.0 | 246.8 | 23.6 | 38.4 | 0.6 | 24.0 | 197.6 | 19.8 |
| P14-2 | F | 28.3 | 192.3 | 25.3 | 43.7 | 1.2 | 23.7 | 160.7 | 20.8 |
| P14-3 | F | 24.0 | 208.0 | 19.3 | 54.8 | 0.7 | 19.3 | 130.5 | 17.7 |
| P14-4 | F | N/A | N/A | N/A | N/A | N/A | 15.0 | 171.2 | 13.2 |
| P14-5 | M | 20.8 | 253.6 | 17.0 | 106.0 | 0.8 | N/A | N/A | N/A |
| P14-6 | M | 21.2 | 213.7 | 19.2 | 47.5 | 0.5 | N/A | N/A | N/A |
| P14-7 | M | 34.8 | 225.7 | 28.8 | 67.8 | 0.7 | N/A | N/A | N/A |
| P21-1 | F | 31.0 | 246.8 | 28.2 | 124.2 | 0.8 | N/A | N/A | N/A |
| P21-2 | F | 67.0 | 158.5 | 44.0 | 41.5 | 2.5 | 58.8 | 154.5 | 26.8 |
| P21-3 | F | 44.7 | 195.8 | 37.8 | 51.3 | 1.2 | 52.8 | 159.7 | 37.7 |
| P21-4 | F | 42.8 | 194.8 | 38.5 | 56.3 | 0.8 | 46.7 | 142.8 | 34.7 |
| P21-5 | F | 29.0 | 162.2 | 25.0 | 52.5 | 0.5 | 27.5 | 158.5 | 22.7 |
| P21-6 | M | 60.2 | 179.2 | 54.8 | 65.8 | 2.7 | 62.2 | 150.8 | 57.5 |
| P21-7 | M | 23.0 | 136.7 | 17.7 | 61.2 | 0.8 | 13.8 | 76.0 | 9.2 |
| P55-1 | F | 215.5 | 54.5 | 50.0 | 62.3 | 1.3 | 43.2 | 136.3 | 28.2 |
| P55-2 | F | 226.8 | 60.2 | 48.2 | 197.4 | 1.0 | N/A | N/A | N/A |
| P55-3 | F | 217.2 | 66.0 | 56.2 | 152.8 | 1.4 | N/A | N/A | N/A |
| P55-4 | M | 267.0 | 66.0 | 55.3 | 121.2 | 2.2 | 54.0 | 171.7 | 41.7 |
| P55-5 | M | 267.3 | 66.8 | 57.0 | 131.2 | 2.8 | 49.7 | 156.7 | 35.7 |

**Supplementary Table 4:** Individualized marker quantification and colocalization with claustrum cells that express cFos for each mouse in the naive and open field (OF) groups. Numbers represent mean cell count per slice, averaged across the anterior, middle, and posterior claustrum. The first column reflects mouse age, where P(x) indicates the postnatal day when tissue was collected. F: female, M: Male.

| Mouse ID | Sex | Group | cFos | Nurr1 | cFos+/Nurr1+ | Tle4 | cFos+/Tle4+ |
| --- | --- | --- | --- | --- | --- | --- | --- |
| P90-1 | F | Naive | 13.5 | 113.7 | 7.7 | 137.2 | 0.7 |
| P90-2 | F | Naive | 21.3 | 444.5 | 14.2 | 74.8 | 0.7 |
| P90-3 | M | Naive | 9.2 | 181.2 | 5.0 | 123.2 | 0.0 |
| P90-4 | M | Naive | 10.0 | 117.0 | 5.3 | 103.2 | 0.3 |
| P90-5 | M | Naive | 0.7 | 124.7 | 0.2 | 114.2 | 0.0 |
| P90-6 | F | OF | 66.0 | 213.2 | 39.0 | 113.3 | 4.2 |
| P90-7 | F | OF | 59.3 | 181.0 | 37.0 | 94.7 | 2.8 |
| P90-8 | F | OF | 51.5 | 228.5 | 28.7 | 171.3 | 1.5 |
| P90-9 | F | OF | 60.2 | 206.8 | 32.0 | 261.0 | 0.3 |
| P90-10 | F | OF | 104.7 | 273.7 | 67.5 | 178.0 | 7.8 |
| P90-11 | F | OF | 63.3 | 211.0 | 48.7 | 149.3 | 3.2 |
| P90-12 | M | OF | 70.3 | 211.0 | 53.0 | 117.7 | 4.7 |
| P90-13 | M | OF | 68.0 | 222.3 | 44.3 | 137.3 | 5.7 |
| P90-14 | M | OF | 49.8 | 177.2 | 24.2 | 84.7 | 2.7 |
| P90-15 | M | OF | 64.7 | 200.3 | 35.0 | 129.0 | 6.8 |

## 8. DECLARATIONS

### 8.1. Ethics approval and consent to participate

All procedures were performed in accordance with the Canadian Council on Animal Care Guidelines and were approved by the University of Alberta Animal Care and Use Committee (AUP-3715).

### 8.2. Consent for publication

Not applicable.

### 8.3. Availability of supporting data and materials

All data are available from the corresponding author upon request.

### 8.6. Authors’ contributions

J.J. and T.S. conceived and designed the project. J.J. supervised the project. T.S. performed most of the stereotaxic injections in adult mice with assistance from V.C., B.A.M. and M.S. All injections in neonatal mice were performed by T.S. All behavioural experiments were performed by G.J.D. T.S. and G.J.D. collected tissue, performed immunohistochemistry and confocal imaging. T.S. and G.J.D. analyzed the data with contributions from J.J. and M.A. The manuscript was written by T.S. with assistance from G.J.D., and edited by J.J. All authors read and approved the final version of the manuscript.

## 8.7. Acknowledgements

We thank cell imaging core at the University of Alberta (the Katz Group Centre and the Cross Cancer Institute) for their support.

